# Integrative analysis of CAM photosynthesis reveals its impact on primary metabolism in *Yucca*

**DOI:** 10.1101/2024.11.01.621533

**Authors:** David Wickell, Richard Field, Karl Weitz, Rosalie Chu, Jesse Trejo, Nikola Tolic, Nathalie Munoz Munoz, Hope Hundley, Emily Savage, Anna Lipzen, Jim Leebens-Mack, Karolina Heyduk

**Affiliations:** Department of Ecology and Evolutionary Biology, University of Connecticut, Storrs, CT, 06269; Department of Plant Biology, University of Georgia, Athens, GA, 30602; USDA-ARS, Athens, GA, 30605; Environmental Molecular Sciences Laboratory, Pacific Northwest National Laboratory, Richland, WA, 99354; Biological Sciences Division, Pacific Northwest National Laboratory, Richland, WA, 99354; US Department of Energy Joint Genome Institute, Lawrence Berkeley National Laboratory, Berkeley, CA, 94720

**Keywords:** Crassulacean Acid Metabolism (CAM), *Yucca*, metabolomics, proteomics, transcriptomics, photorespiration, nitrogen metabolism, primary metabolism

## Abstract

Crassulacean Acid Metabolism (CAM) is an adaptation that temporally separates carbon uptake at night from photosynthesis during the day. CAM has evolved repeatedly across vascular plants, as its emergence may depend on simple regulatory changes to deeply conserved metabolic pathways. Modern CAM research relies heavily on interpretation of transcriptomic data, though regulation occurs at multiple levels following transcription. Additionally, while most research to date has focused on a handful of genes and metabolites in the core CAM pathway, the co-option of conserved regulatory and functional genes is bound to have wide ranging effects on other aspects of primary metabolism. In this study, we integrate transcriptomic, proteomic, and metabolomic data to compare primary metabolism between the CAM species *Yucca aloifolia* and closely related C_3_ species, *Y. filamentosa*. We observe minimal correlation between protein abundance and mRNA expression, suggesting significant post-transcriptional regulation in CAM species. We also find evidence of shifts in gene expression and metabolite accumulation outside of the central CAM pathway suggesting that the shift to CAM has cascading effects across primary metabolism, especially nitrogen metabolism. Our findings provide insights into the metabolic shifts associated with CAM evolution and highlight the complexity of its regulation at multiple biological levels.

## Introduction

Crassulacean Acid Metabolism (CAM) is a variation on the more common C_3_ metabolism that allows plants to temporally separate carbon uptake and storage from photosynthesis. Briefly, in C_3_ plants, uptake of atmospheric CO_2_ and carbon fixation by Rubisco co-occur during the day. In CAM plants, however, CO_2_ is taken up at night and stored in the vacuole as malate before being liberated and fed into the Calvin cycle during the day (Fig. 1). CAM photosynthesis allows terrestrial plants to keep their stomata closed during the hottest part of the day, greatly reducing transpirational water loss. This, in combination with reduced rates of photorespiration due to the enrichment of CO_2_ in photosynthetic tissues during the day, results in dramatic improvement to water use efficiency in terrestrial CAM lineages relative to C_3_ plants (Winter and Smith 2022).

**Figure 1:**
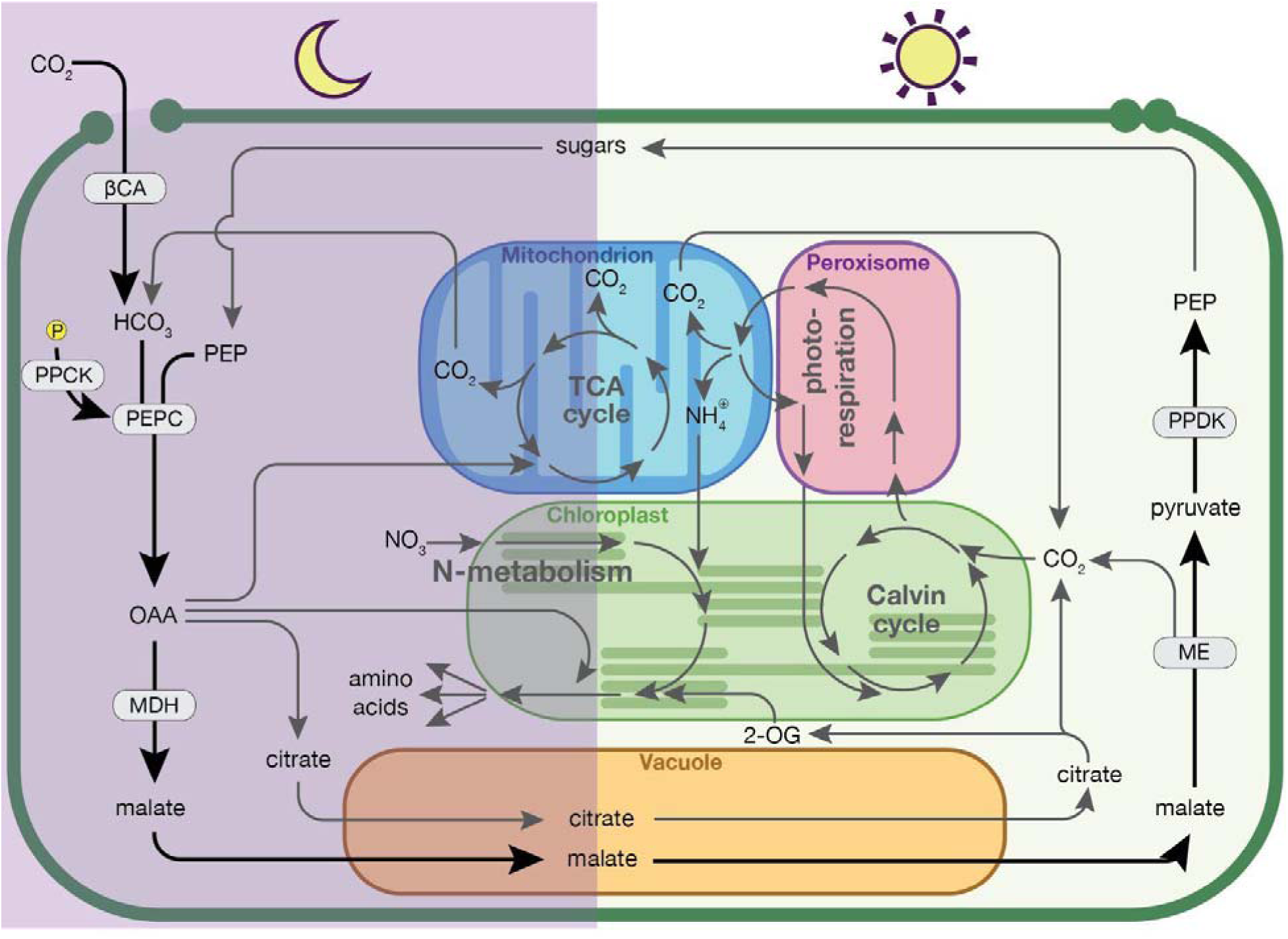
Flow chart of the core CAM pathway. (black arrows) and its interactions with other primary metabolic pathways (gray arrows). Alternatively, malate can be decarboxylated by MDH and PEPCK during the day (not shown) and conversion of citrate to 2-OG may occur in the cytosol (as shown) or mitochondrion (not shown). Abbreviations: 2-OG, 2-oxoglutarate; βCA, beta carbonic anhydrase; MDH, malate dehydrogenase; ME, malic enzyme; OAA, oxaloacetate; PEP, phosphoenolpyruvate; PEPC, phosphoenolpyruvate carboxylase; PPDK, pyruvate-phosphate dikinase.

CAM is distinct from other carbon concentrating mechanisms in that it temporally separates CO₂ uptake from Rubisco-mediated fixation, whereas C₄ photosynthesis and pyrenoids achieve this separation spatially, relying on specialized anatomy (Sage et al. 2012) or subcellular compartments (Robison et al. 2025), respectively. Accordingly, CAM physiology appears to arise from regulatory modifications to deeply conserved pathways common to all green photosynthetic organisms. Still, there is some debate as to whether the emergence of CAM requires complex regulatory rewiring across the genome (Wai et al. 2019), if it is the product of minimal changes resulting in increased metabolic flux through existing C_3_ pathways (Bräutigam et al. 2017), or some combination of the two. Despite the potential complexity of such fundamental alterations to primary metabolism, CAM has evolved over 60 times across the plant phylogeny (Gilman et al. 2023). It is found in every major lineage of vascular plants, in diverse taxa occupying an equally diverse range of habitats from deserts to lakes to rainforest canopies. While the central components of the CAM pathway are present in highly divergent taxa, studies have found a great deal of lineage specific variation as well. CAM plants can vary in their mechanism of day-time decarboxylation (Ramachandra Reddy and Rama Das 1978), gene copy recruitment (Wickell et al. 2021; Heyduk et al. 2022), the role of circadian regulation (Wickell et al. 2021; Chen et al. 2020; Wai et al. 2019), and citrate accumulation (Lüttge 1988; Franco et al. 1992; Freschi et al. 2010). This high degree of interspecific variation in the details of CAM makes comparison of closely related CAM and C_3_ species essential to our understanding of a complex syndrome that has evolved repeatedly in a diverse array of genomic backgrounds.

To date, most genome-enabled research on CAM species has focused on the central carboxylation mechanisms that fix CO_2_ into malate for storage at night and decarboxylation processes that release CO_2_ during the day (e.g. Wickell et al. 2021; Ming et al. 2015; Abraham et al. 2016). These core CAM pathways involve genes, proteins, and metabolites that are also central components of primary metabolism in C_3_ plants where they replenish intermediates in the citric acid cycle. As such, alterations to their expression and abundance are expected to have complex downstream effects on pathways outside of CAM. A notable example of the effects of CAM outside of the canonical pathway is the night time accumulation of citrate in many CAM plants, resembling that of malate (Franco et al. 1992; Freschi et al. 2010; Miszalski et al. 2013; Kornas et al. 2009). Authors have suggested multiple potential explanations that integrate citrate into the existing CAM framework including recapture of respired CO_2_ (Franco et al. 1992), control of redox potential (Kornas et al. 2009), and mitigation of photoinhibition and photorespiration via efficient concentration of CO_2_ around Rubisco (Kornas et al. 2009; Miszalski et al. 2013). However, unlike malate, citrate synthesis is not associated with a net carbon gain (Lüttge 1988). Thus, despite its frequent association with CAM, the precise role of nocturnal citrate accumulation and how frequently it occurs across CAM lineages remain understudied and ultimately unresolved.

Organic acid accumulation, including malate and citrate, has been previously shown to be impacted by both the relative abundance and source of nitrogen and several studies have sought to characterize a mechanistic relationship between CAM and nitrogen metabolism (Santos and Salema 1991; Pereira et al. 2018; Gonçalves and Mercier 2021; Pereira and Cushman 2019). Specifically, researchers have found that some species including *Guzmania monostachia* and several *Kalanchoë* species exhibit increased CAM expression and nocturnal acidification under nitrogen deficiency, further underscoring connections between carbon and nitrogen metabolism (Rodrigues et al. 2014; Winter et al. 1982; Ota 1988; Nobel 1983). However, few studies have incorporated gene expression data and detailed comparisons of N-metabolism between closely related CAM and C_3_ lineages are rare. Similarly, few papers have investigated the photorespiratory pathway in CAM lineages (Duarte and Lüttge 2007; Lüttge 2011) despite it being widely accepted that the reduction of photorespiration is one of the primary factors driving the convergent evolution of CAM in vascular plants.

The Agavoideae has emerged as an important model group for studying the evolutionary origins of CAM due to its high degree of photosynthetic diversity, containing C_3_, CAM, and physiologically intermediate “C_3_ + CAM” species (Heyduk et al. 2023). In particular, the genus *Yucca* within the Agavoideae is well suited to comparative analyses of CAM evolution as it comprises both C_3_ and CAM species with a single origin of CAM inferred roughly 6 mya in section Yucca (Heyduk, McKain, et al. 2016; McKain et al. 2016). The presence of closely related species that differ in photosynthetic pathway allows researchers to investigate CAM through comparative analyses while controlling for the confounding effects of shared ancestry. As a result, considerable research in *Yucca* has investigated the genomic (Heyduk et al. 2019, 2022), physiological (Heyduk, Burrell, et al. 2016; Heyduk et al. 2020), and anatomical (Heyduk et al. 2020; Heyduk, McKain, et al. 2016) changes associated with the transition to CAM. A recent study found evidence of shared expression patterns of core CAM genes between closely related C_3_ and CAM species (Heyduk et al. 2019). Similarly, ancestral state reconstruction in Agavoideae indicated that key features of CAM anatomy such as leaf thickness, cell succulence, and 3D venation may have evolved prior to strong CAM (Heyduk, McKain, et al. 2016). These studies seem to corroborate work suggesting that the evolution of CAM may merely require an increase in flux through existing carboxylation pathways (Bräutigam et al. 2017). Even so, Heyduk et al. (2019) also found widespread differences in gene expression within and outside of the central CAM pathway alongside differential abundance of several primary metabolites between CAM and C_3_ species, indicative of potentially complex downstream effects of CAM across primary metabolism.

Characterizing the wide-ranging effects of CAM on primary metabolism requires simultaneous comparative analysis of multiple pathways and molecules. While this would ideally integrate functional genetic analysis, these types of studies are still in their infancy in CAM plants; *Kalanchoë* is the only CAM lineage where knockdown studies have confirmed genetic function (Dever et al. 2015; Agarie et al. 2020; Ferrari et al. 2020; Boxall et al. 2020). As such, comparative multi-omics approaches potentially offer a more accessible way to investigate modification to primary metabolism in a greater diversity of CAM species. In this study we integrated transcriptomic, proteomic, and metabolomic time series data for *Yucca aloifolia* (CAM) and *Y. filamentosa* (C_3_) to investigate shifts in primary metabolism associated with the transition from C_3_ to CAM. Comparison of transcriptomic and proteomic datasets showed little correlation between fluctuations in transcript and protein abundance over time, suggesting that post-translational modification and activation/inhibition by other proteins and metabolites might play a significant role in regulation of CAM following translation of mRNA into proteins. We also found alterations to protein abundance, gene expression, and metabolite abundance related to citrate metabolism and nitrogen metabolism between *Y. aloifolia* (CAM) and *Y. filamentosa* (C_3_). However, despite interspecies differences in the abundance of a few metabolites associated with photorespiration, there were few clear differences in protein abundance or gene expression between the two species. This potentially suggests a diminished role for alterations to the photorespiratory pathway in the evolution of CAM in *Yucca*. Our findings underscore the utility of comprehensive multi-omics analyses in clarifying complex metabolic shifts associated with CAM, providing valuable insights into gene regulation, and generating testable hypotheses that pave the way for further research of CAM evolution across vascular plants.

## Materials and Methods

### Physiology

Six mature *Y. aloifolia* plants, each a different genotype, were selected from a collection of wild-collected plants at the University of Georgia (UGA) Plant Biology greenhouse. Six mature *Y. filamentosa* plants, also each a different genotype, were obtained either from the UGA collection or were purchased from landscape nurseries. Plants obtained from nurseries were re-potted in 5-gallon containers with a 50/50 mixture of standard potting soil and sand (the same mixture in which the UGA collection plants were grown and maintained) and allowed to acclimate to the greenhouse for at least 4 weeks. Plants were introduced into a growth chamber at the UGA Plant Biology greenhouse in randomized positions one week before the onset of sampling to allow entrainment to the light regime and were watered daily. Growth chamber conditions were programmed with the following conditions: the light period commenced at 7:00 am with relative humidity of 40%, ambient temperature at 30° C, and photosynthetically active radiation (PAR) approximately 270 μmol m^-2^ s^−1^ measured at leaf level. The dark period commenced at 7:00 pm with relative humidity of 40%, ambient temperature at 17° C, and PAR 0 μmol m^−2^ s^−1^.

Gas exchange and leaf tissue sampling for the well-watered treatment commenced on December 03, 2021. Beginning one hour after the onset of the light period (“Zeitgeber time 1”, ZT1), instantaneous gas exchange measurements were obtained with a LiCOR-6400XT portable photosynthesis system on mid-leaf sections of mature leaves. LiCOR chamber conditions were set to match ambient conditions in the growth chamber, including temperature, light, and humidity; the fan speed was set to maximum and flow was set at 500 umol s^-1^. Gas exchange measurements were repeated on the same leaf, and a random leaf was sampled for each plant, in the same order every four hours for the next five time points (ZT5, ZT9, ZT13, ZT17, ZT21). Immediately following gas exchange measurements, a randomly selected, mature leaf was excised and snap frozen in liquid N_2_ and stored at -80° C. Soil moisture was measured at the end of the 24-hour sampling period with a digital soil moisture probe inserted into the top ∼3 inches of soil for each plant. Watering was then withheld for the remainder of the experiment for “drought” treated plants. Soil moisture levels were periodically checked over the next 18 days. At day 13, it was observed that the *Y. aloifolia* species’ soil was drying down faster than that of *Y. filamentosa*. To maintain comparable soil conditions, each fast-drying plant was top-watered with 500 mL water, bringing those plants back to similar soil water content as the others. By day 18 (December 20^th^, 2021), the average soil moisture for each species was below 5% and gas exchange and leaf tissue sampling were repeated in the same manner as the well-watered samples.

### RNA-sequencing and analysis

Whole leaf tissue for each sample was ground to a powder in a mortar and pestle under liquid N_2_, transferred to a 15 mL Falcon tube and stored in a -80° C freezer. RNA extraction for each sample from an aliquot of approximately 100 mg of powdered leaf tissue was carried out with the Qiagen Plant RNeasy kit with DNase I on-column digestion according to the Qiagen standard protocol. RNA quantity was measured with a Qubit fluorometer, and RNA integrity was measured on an Agilent 2100 Bioanalyzer with an RNA Nano 6000 kit. RNA samples were shipped on dry ice to the DOE’s Joint Genome Institute (JGI) labs in Berkeley, CA for cDNA library construction and paired-end 150 bp Illumina sequencing on an Illumina NovaSeq sequencer. All but five libraries produced deep sequence data (total PE150 libraries = 139, median fragment count = 28,066,771, mean Q30 = 0.95).

### RNA-seq read quantification

RNA sequence data was downloaded from the JGI server onto the Georgia Advanced Computing Research Center computing cluster. For each species, gene annotations were used from the draft genomes on Phytozome (Yucca aloifolia Ya24Inoko v2.1 DOE-JGI, https://phytozome.jgi.doe.gov/info/YaloifoliaYa24Inoko_v2_1 and Yucca filamentosaC3 pri v2.1 DOE-JGI, https://phytozome.jgi.doe.gov/info/YfilamentosaC3pri_v2_1) (pre-publication permission from JGI was obtained for *Yucca* annotation use). Reads for each species were pseudo-aligned to their respective primary transcript nucleotide fasta sequences and quantification was calculated with *kallisto* v0.48.0 (Bray et al., 2016) with default parameters. Raw read counts were imported into expression matrices in R with *tximport* v1.14.2 (Soneson et al., 2016).

### Inference of homology with Orthofinder

Homology was assessed with Orthofinder2 (Emms and Kelly 2019) using CDS sequences from the *Y. aloifolia* and *Y. filamentosa* genome assemblies from Phytozome along with genomic CDS sequences from the following taxa: *Acorus americanus* (Acorus americanus v1.1, DOE-JGI, http://phytozome-next.jgi.doe.gov/), *Ananas comosus* (Ming et al. 2015), *Asparagus officinalis* (Harkess et al. 2017), *Agave tequiliana* (Agave tequilana var. Weber’s Blue v2.1 DOE-JGI, https://phytozome-next.jgi.doe.gov/info/Atequilanavar_WebersBlueHAP1_v2_1), *Arabidopsis thaliana* (Cheng et al. 2017), *Amborella trichopoda (Amborella Genome Project 2013)*, *Brachypodium distachyon* (International Brachypodium Initiative 2010), Kalanchoe fedtschenkoi (Yang et al. 2017), *Oryza sativa* (Ouyang et al. 2007), *Sorghum bicolor* (McCormick et al. 2018), *Solanum lycopersicum* (Tomato Genome Consortium 2012), and *Zea mays* (Hufford et al. 2021). CDS sequences were translated to amino acids prior to running Orthofinder using the default settings.

### Time-structured expression profiling of transcriptomic data

To assess differential gene expression patterns due to watering regimes over the 24-hour sampling period, differential expression analysis taking time and treatments as variables was carried out in the R software package *maSigPro* v.1.58.0 (Nueda et al., 2014). For each species, trimmed mean of *M* values (TMM) were calculated and used to normalize read counts within species, and counts per-million (CPM) were computed from the normalized libraries in the R software package *edgeR* v3.28.0 (Robinson et al. 2009). Genes with less than 0.5 CPM across all time points were removed. Outlier libraries were checked with multidimensional scaling plots. Two and six libraries were removed from *Y. aloifolia* and *Y. filamentosa*, respectively. Genes were tested for significant time structure with functions *p.vector (counts=TRUE)*, and the *T.fit(step method = “backward”, alpha = 0.05)*. Genes with high variability between replicates were classified as overly influential by *maSigPro*, removed, and the analyses were rerun on the filtered data set. Genes with significant treatment effects across time were called with *sig.siggenes*(*rsq* = 0.6). To sort genes into clusters based on time-structured expression similarity, *maSigPro* requires the user to input the expected numbers of clusters (*k*). To estimate *k,* within-group sums of squares were calculated and plotted so the *k* value at which marginal reductions in the sums of squares can be determined. *k*=6 was chosen for both species and genes were sorted into clusters using *see.genes(cluster.method="hclust")*.

### Metabolite extraction and protein isolation

2.0 ml tubes (Safe-Lock, Eppendorf) were filled with leaf material to approximately ¼ capacity, covered with tube membranes, and samples were lyophilized overnight. Lyophilized leaves were then homogenized in a Tissue Lyser II (Qiagen) using tungsten beads, specifically two-3mm beads and one-5mm in each tube. Samples were lysed for 2 min at a frequency of 23 Hz until ground to a powder. Metabolites from 20 mg of the ground leaves were extracted with 800 µl of a 4:1 methanol: water mix, and each sample spiked with 50 µl of a 330 µg/ml solution of glycine d5 (Cambridge Isotopes). Samples were then placed in a thermomixer and shaken at 1200 rpm and 21°C for 1 hr. Following centrifugation at 5000 x g, supernatant containing the metabolite fraction was removed and 10% of the total metabolite extract was transferred to a glass vial to be dried under vacuum for subsequent metabolomic analysis (described below). After removing the metabolite fraction, protein pellets were washed twice by adding 500 µl 100% MeOH, vortexing for 15 seconds, and centrifuging at 10000 x g and 21°C for 10 min. The resulting pellet was dried in a fume hood until remaining MeOH had completely evaporated and stored at -80°C until protein digestion.

Protein pellets were resuspended in 120 µl of an aqueous solution of 8 M urea and 50 mM HEPES by vortexing and sonication for 30 sec. Following resuspension, 6.3 μL 1 M aqueous solution of DTT was added to each sample and samples were placed in a thermomixer and shaken at 500 rpm and 37°C for 30 min for reduction. Next, 40 μL 0.5 M aqueous solution of iodoacetamide was added to each sample and samples were placed in a thermomixer and shaken for another hour at 500 rpm and 37°C for alkylation. Samples were then diluted with 800 μL 50 mM aqueous solution of HEPES prior to digestion with trypsin (1:50 w/w trypsin:protein) for 3 hrs at 500 rpm and 37°C. Sample clean-up was performed using C18 SPE media and samples were concentrated to ∼100 μL and peptide concentration was determined via BCA assay. Finally, 30 μg of each sample were transferred to a new 1.5 mL tube, dried down in a speedvac and stored at -80°C for TMT labeling.

### TMT isobaric tag labeling

Samples were designed into 12 TMT sets (6 per plant) including two pools, one consisting of a mix of the well-watered plants and one of all the drought plants to ensure correlation across sets. Each set included 1 replicate of each condition and blanks to establish a base background noise signal and then each set completely randomized. Each sample was diluted in 40 µl 500 mM HEPES, pH 8.5 and labeled using amine-reactive Tandem Mass Tag Isobaric Mass Tagging Kits (Thermo Scientific, Rockford, IL) according to the manufacturer’s instructions. Briefly, 250 μL of anhydrous acetonitrile was added to each 5 mg reagent, vortexed, and allowed to dissolve for 5 mins with occasional vortexing. Reagents (5 µl) were then added to each sample and incubated for 1 hour at room temperature with shaking at 400 rpm. Each sample was then diluted with 30 µl 20% acetonitrile. A portion from each sample was collected as a premix to run on a mass spectrometer to ensure complete labeling. The samples were frozen at -80°C until the results showed efficient labeling. At that point the frozen samples were thawed and the reaction was quenched by adding 8 μL of 5 % hydroxylamine to the sample with incubation for 15 mins at room temperature with shaking at 400 rpm. The samples within each set were then combined and completely dried in the speedvac. Each of the samples were then resuspended in 0.1% trifluoroacetic acid solution and cleaned using Discovery C18 50 mg/1 mL solid phase extraction tubes as described above and once again assayed with bicinchoninic acid to determine the final peptide concentration and vialed for microfractionation.

### Microfractionation

Peptide mixtures were separated by high resolution reversed phase UPLC using a nanoACQUITY UPLC® system (Waters Corporation, Milford, MA) equipped with an autosampler. Capillary columns, 200 µm i.d. × 65 cm long, were packed with 300Å 3.0 µm p.s. Jupiter C18 bonded particles (Phenomenex, Torrence, CA). Separations were performed at a flow rate of 2.2 µL/min on binary pump systems, using 10 mM ammonium formate (pH 7.5) as mobile phase A and 100% acetonitrile as mobile phase B. 20 µL of TMT labeled peptide mixtures (0.5 µg/µL) were loaded onto the column and separated using a binary gradient of 1 % B for 35 min, 1-10 % B in 2 min, 10-15 % B in 5 min, and 15-25 % B in 35 min, 25-32 % B in 25min, 32-40 % B in 13 min, 40-90 % B in 43 min, held at 90 % B for 2 min (column washed and equilibrated from 90-50 % B in 2 min, 50-95% B in 2 min, held at 95 % B for 2 min and 95-1 % in 4 min). The capillary column eluent was automatically deposited every minute into 12x32 mm polypropylene vials (Waters Corporation, Milford, MA) starting at minute 60 and ending at minute 170, over the course of the 180 min LC run. Fractions were concatenated by collecting fractions from vial 1 to vial 24 and then returning to vial 1 back to vial 24 (etc.). Prior to peptide fraction collection, 100 µL of water was added to each vial to aid the addition of small droplets into the vial. Each vial was completely dried in a vacuum concentrator (Labconco, Kansas City, MO), reconstituted in 15 µL 2 % acetonitrile, 0.1 % formic acid and stored at -20°C until LC-MS/MS analysis.

### TMT quantitation of isolated peptides

A Waters nano-Acquity dual pumping UPLC system (Milford, MA) was configured for on-line trapping of a 5 µL injection at 5 µL/min for 5 min with reverse-flow elution onto the analytical column at 200 nL/min using mobile phase A. The analytical column was slurry packed in-house using Waters BEH C18 1.7 um particles (Waters Chromatography, Drinagh, Ireland) into a 25 cm PicoFrit 360um od x 75um id column (New Objective, Littleton, MA) and held at 45C during use by a 15 cm AgileSleeve flexible capillary heater (Analytical Sales and Services, Inc., Flanders, NJ). The trapping column was slurry packed in-house using Jupiter 5um C18 particles (Phenomenex, Torrence, CA) into a 4cm long piece of 360um od x 150um id fused silica (Polymicro Technologies, Phoenix, AZ) and solgel fritted at both ends. Mobile phases consisted of (A) 0.1% formic acid in water and (B) 0.1% formic acid in acetonitrile with the following gradient profile (min, %B): 0, 1; 10, 8; 105, 25; 115, 35; 120, 75; 123, 95; 129, 95; 130, 50; 132, 95; 138, 95; 140, 1.

MS analysis of 24 fractions per plex set was performed using a Orbitrap Fusion Lumos Tribrid MS (ThermoFisher Scientific) outfitted with a homemade nano-electrospray ionization interface. Electrospray emitters were homemade using 150 μm o.d. × 20 μm i.d. chemically etched fused silica. The ion transfer tube temperature and spray voltage were 250°C and 2.2 kV, respectively. Peptides were separated on an in-house prepared 40 cm x 75 um 1.7-um Waters BEH C18 column, mobile phases consisted of 0.1% formic acid in water (MP-A) and 0.1% formic acid in acetonitrile (MP-B). Data were collected for 120 min following a 10 min delay after completion of sample trapping and start of LC gradient.

FT-MS spectra were acquired from 350 to 1800 m/z at a resolution of 60 k (AGC target 4e5) and the FT-HCD-MS/MS spectra were acquired in data-dependent mode for ions with detected charge states 2-6 and minimum intensity of 2.5e3 using a normalized collision energy of 30. Other MS/MS parameters were as follows - isolation window was set to 0.7 m/z at a resolution of 50 k, AGC target 1e5, and dynamic exclusion time was 45 s.

Acquired raw spectra were processed using automated MSGF+ software (Kim et al. 2008) in target/decoy mode with 20 ppm parent ion mass tolerance, partial tryptic rule with oxidation of methionine (+15.9949) as dynamic modification and static modifications carbamidomethylation of cysteine (+57.0215) and TMTpro modification of lysine on peptide N termini (+304.2071). Each set of samples was matched with equal-contribonding protein collections for *Y. aloifolia* (43,466 protein sequences) and *Y. filamentosa* (45,266 protein sequences). Common contaminant proteins from trypsin and keratins (16 protein sequences) were considered in each search.

Top peptide-spectral matches (PSM) passing 1% FDR cut-off established for the whole datasets of *Y. aloifolia* and *Y. filamentosa* were used for subsequent TMT quantification. MASIC (Monroe et al. 2008) software was used to pull abundances of the sample reporter ions for all PSMs. Measured reporter ion abundances were log transformed to remove skewness of abundance distribution and normalized using mean central tendency normalization. Peptide abundances from all fractions were summed prior to normalization, normalization was performed using InfernoRDN (Polpitiya et al. 2008) software. Protein level roll-up of normalized peptide abundances was normalized again using the same strategy and software.

### Proteomic abundance analysis

Normalized protein abundance data was filtered to remove proteins supported by less than two peptide matches as well as proteins that were missing abundance data for more than ⅓ of samples. After filtering, abundance values were converted to z-scores in R to facilitate removal of outlier data points, identification of proteins that fluctuate in abundance throughout the day (a.k.a. “cycling” proteins), and high level comparisons between species and timepoints. For each gene and treatment, individual abundance values more than two standard deviations above or below the next closest abundance value at a given time point were considered outliers and removed prior to identifying cycling proteins.

Cycling proteins were identified and peak mean expression was calculated using a custom R script. To identify cycling proteins, filtered, z-scaled abundance data was divided into drought and well-watered conditions prior to fitting polynomial functions for linear, quadratic, cubic, and quartic models. Model fits were compared using ANOVA, and the best-fitting model was defined as the polynomial degree associated with the smallest ANOVA p-value. Proteins were considered cycling if any higher-order model provided a significantly better fit than the linear model (*p* < 0.05). R scripts used for z-scaling, outlier removal, and identification of cycling proteins are available on GitHub at https://github.com/dawickell/Yucca-omics.

### Untargeted metabolomic quantitation and analysis

Dried metabolite extracts were chemically derivatized as previously reported (Kim et al. 2015). Briefly, the extracts were derivatized by methoxyamination and trimethylsilyation (TMS), then the samples were analyzed by GC-MS. Samples were acquired in an Agilent GC 7890A using a HP-5MS column (30 m × 0.25 mm × 0.25 μm; Agilent Technologies, Santa Clara, CA) coupled with a single quadrupole MSD 5975C (Agilent Technologies). One microliter of sample was injected into a splitless port at constant temperature of 250°C. The GC temperature gradient started at 60 °C, with a hold of temperature for 1 minute after injection, followed by increase to 325 °C at a rate of 10 °C/minute and a 10-minute hold at this temperature. A fatty acid methyl ester standard mix (C8-28) (Sigma-Aldrich) was analyzed in parallel as standard for retention time calibration.

GC-MS raw data files were processed using the Metabolite Detector software (Hiller et al. 2009). Retention indices (RI) of detected metabolites were calculated based on the analysis of a FAMEs mixture, followed by their chromatographic alignment across all analyses after deconvolution. Metabolites were initially identified by matching experimental spectra to a PNNL augmented version of Agilent GC-MS metabolomics Library, containing spectra and validated retention indices for over 1200 metabolites. Then, the unknown peaks were additionally matched with the NIST20/Wiley11 GC-MS library. All metabolite identification and quantification ions were validated and confirmed to reduce deconvolution errors during automated data-processing and to eliminate false identifications. Peak areas for identified metabolites and unknowns were normalized to the peak area of the internal standard glycine d5, reported and used for further statistical analysis.

### Metabolomic abundance analysis

Time-structured metabolites were identified using polynomial regression and nested ANOVA. Similar to our proteomic analysis, metabolite abundance data was divided by species and condition prior to fitting linear, quadratic, cubic, and quartic polynomial models as a function of time. Model fits were compared using ANOVA, and the best-fitting model was defined as the polynomial degree associated with the smallest ANOVA p-value. Metabolite time courses were considered cycling if any higher-order model provided a significantly better fit than a linear model (p ≥ 0.05), otherwise a linear model was retained.

To test for differences in time-structured abundance between species within a condition, z-scaled abundance data were fit using the maximum polynomial degree selected for that condition. A reduced model including time alone was compared to a full model including a species-by-time interaction using ANOVA. Similarly, to test for differences between conditions within a species, raw abundance values were modeled using the maximum degree selected for that species, and reduced and full condition-by-time interaction models were compared by ANOVA. All scripts used for z-scaling, outlier removal, polynomial model fitting, and statistical testing are available upon request.

### Statistical methods

All analyses were conducted using independent biological replicates. Physiological, transcriptomic, proteomic, and metabolomic data were collected from six biological replicates per species, equal-contribonding to distinct genotypes, sampled under both well-watered and drought treatments, every 4 h over a 24 h period. Gas exchange measurements represent repeated measures on the same individuals over time, while transcriptomic, proteomic, and metabolomic measurements were obtained from independently harvested leaf tissue taken from the same individuals over time.

Time-series data were analyzed using regression-based approaches appropriate for diel sampling designs. Transcriptomic time-structured expression was assessed using multiple regression models implemented in maSigPro <u>(Nueda et al. 2014)</u>, while proteomic and metabolomic time courses were evaluated using polynomial regression and nested analysis of variance (ANOVA) to compare linear and higher-order temporal models. Interaction terms were used to test for differences in temporal patterns between species and between watering treatments. Detailed descriptions of normalization, filtering, model fitting, and significance criteria can be found in the equal-contribonding Methods subsections.

All statistical analyses were performed in R <u>(R Core Team 2021)</u>, and figures were generated using ggplot2 v4.0.0 <u>(Wickham 2011)</u>. Custom analysis scripts are available at https://github.com/dawickell/Yucca-omics

## Results

### Broad comparison of -omics datasets

Gas exchange data (Supplementary Fig. S1) and malate accumulation differences (Fig. 3c) between species support prior evidence that *Y. aloifolia* and *Y. filamentosa* are CAM and C_3_, respectively (Heyduk, Burrell, et al. 2016). Data also corroborated prior work that demonstrated *Y. aloifolia*’s reliance on CAM slightly decreases under (Heyduk et al. 2019). Transcriptomic analysis of *Y. aloifolia* and *Y. filamentosa* found that 83.4% (34,663/41,577) of *Y. aloifolia* genes and 50.4% (22,195/43,996) of *Y. filamentosa* genes exhibited time-structured mRNA expression in well-watered and/or drought treatments (Fig. 2, Supplementary Tables 2 and 3). When analysis was restricted to genes that cycle in both treatments, 54% (22,523/41,577) and 28% (12,357/43,996) of genes cycled in *Y. aloifolia* and *Y. filamentosa*, respectively (Fig. 2, Supplementary Fig. S2, Supplementary Tables 2 and 3).

**Figure 2.**
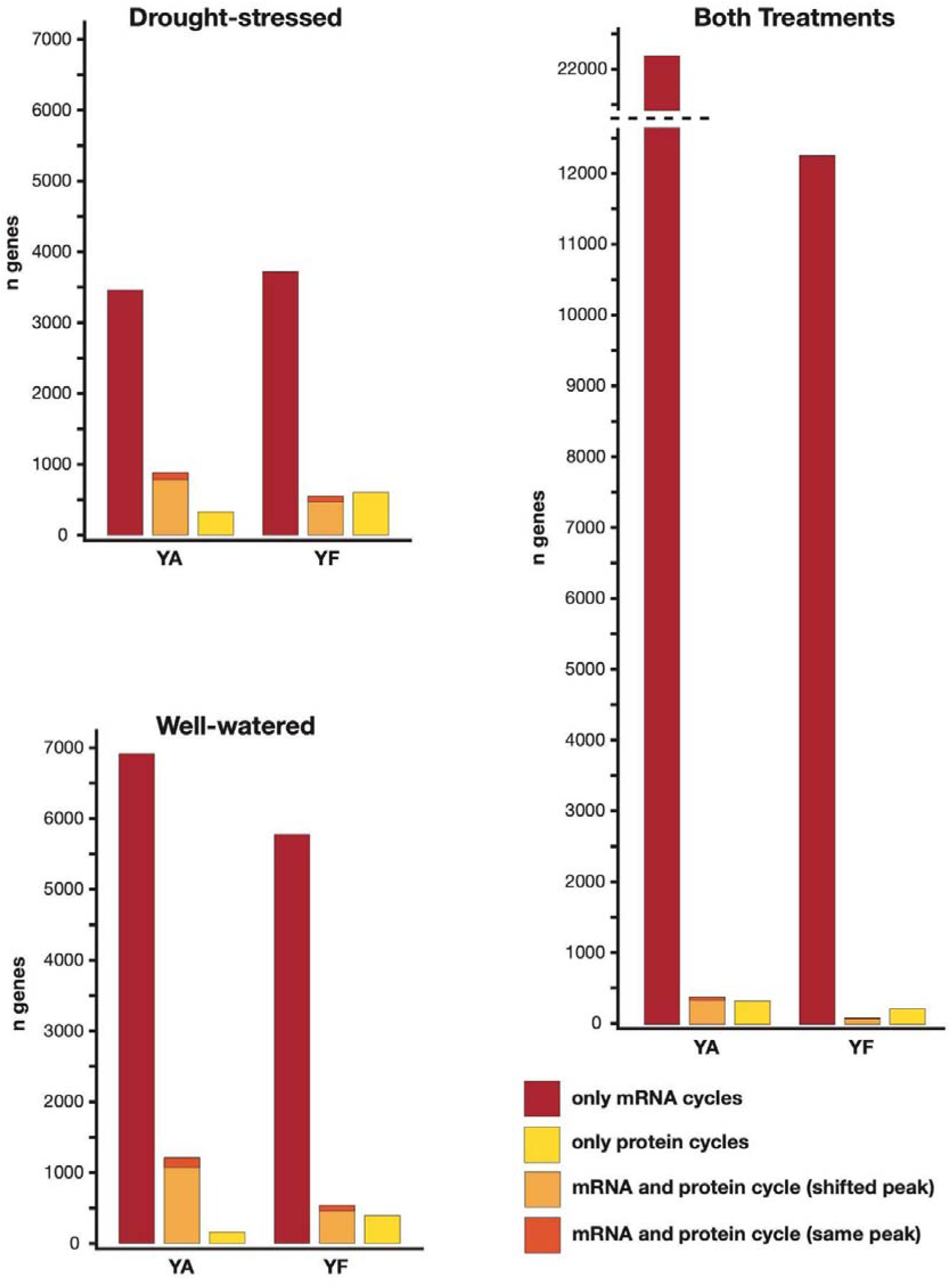
Overlap in temporal patterns of protein abundance and mRNA expression in *Yucca*. Barplots show the total number of genes that exhibit cycling expression and or protein abundance for *Y. aloifolia* (YA) and *Y. filamentosa* (YF). Proteins with abundance peaks shifted ≥4 h relative to its associated mRNA transcript are designated as exhibiting shifted peaks.

**Figure 3:**
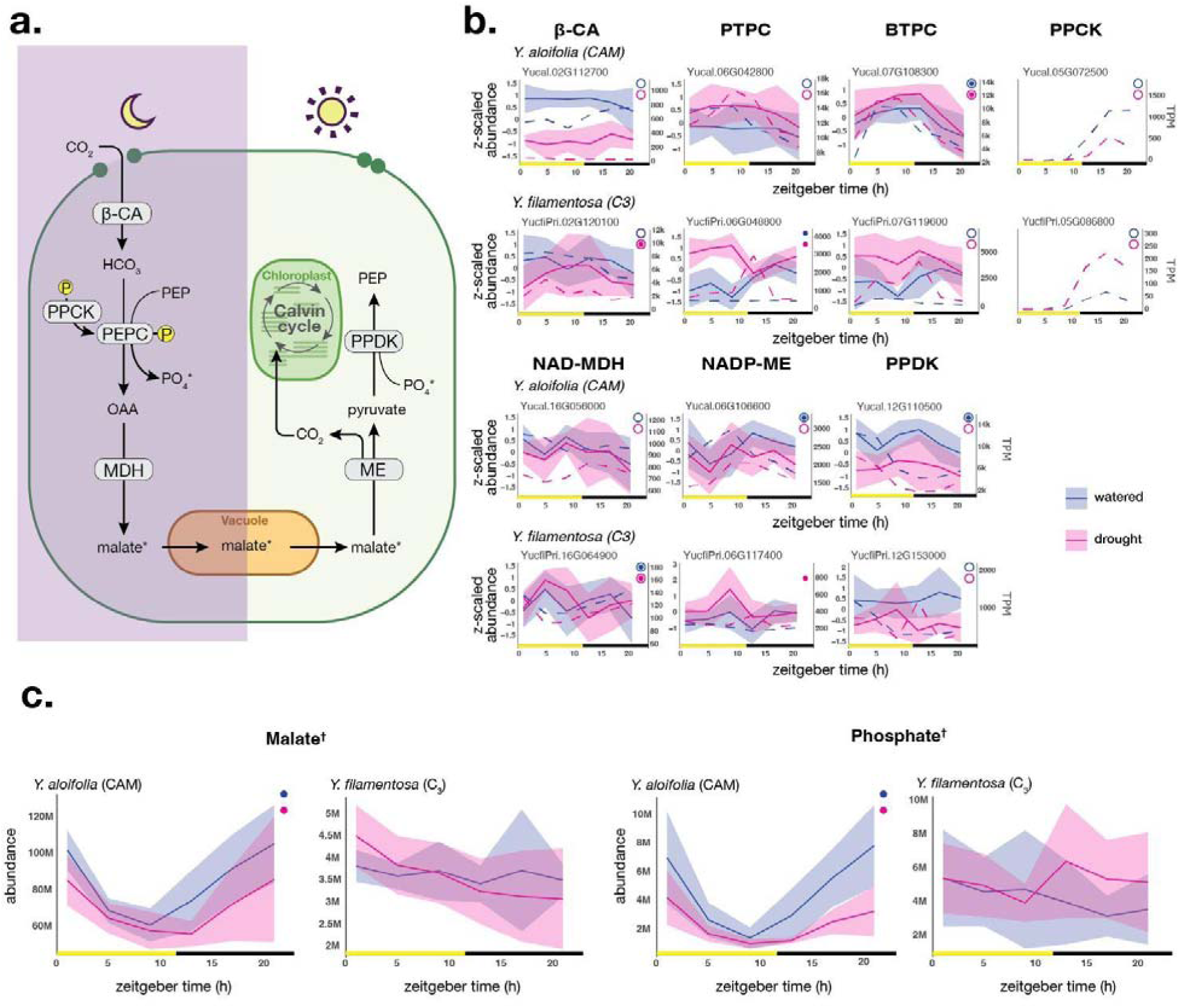
Comparison of CAM photosynthesis in *Y. aloifolia* and *Y. filamentosa.* **(a.)** Flow-chart of core reactions of the CAM pathway in *Y.aloifolia*. **(b.)** Plots of z-scaled mean protein abundance (solid line, SD represented by shaded area) with mean expression in TPM overlaid (dashed line) for comparison. Filled and unfilled circles indicate significantly (*p* < 0.05) time-structured protein abundance and gene expression, respectively. **(c.)** Plots of abundance over time for malate and phosphate, (indicated in flow-chart by an ‘*’) shaded areas represent the standard deviation. Filled circles indicate significantly (*p* < 0.05) time-structured metabolite abundance and daggers (†) indicate metabolites that are significantly (*p* < 0.05) different between species. Abbreviations: b-CA, beta-carbonic anhydrase; PEP, phosphoenolpyruvate; PPCK, phosphoenolpyruvate carboxylase kinase; PEPC, phosphoenolpyruvate carboxylase; PTPC, plant-type phosphoenolpyruvate carboxylase; BTPC, bacterial-type phosphoenolpyruvate carboxylase; OAA, oxaloacetate; NAD-MDH, NAD dependent malate dehydrogenase; NADP-ME, NADP dependent malic enzyme; PPDK, pyruvate phosphate dikinase.

Proteomic analysis of leaf tissue recovered abundance data for far fewer genes than transcriptomic analysis. Abundance was recorded for 6,354 proteins in *Y. aloifolia* and 6,859 proteins in *Y. filamentosa*, representing 15.2% and 15.6% of annotated genes, respectively. Also in contrast to analyses of transcriptomic data, relatively few proteins exhibited time-structured abundance patterns (Fig. 2, Supplementary Fig. S2, Supplementary Tables 4 and 5). In proteins, linear regression identified 50.5% (3,214/6,354) of proteins in *Y. aloifolia* and only 35% (2,400/6,859) of proteins in *Y. filamentosa* as cycling over time in at least one treatment. When the analysis was restricted to genes that cycle in both treatments, far fewer proteins were identified with 9.8% (622/6,354) and 4.5% (310/6,859) proteins cycling in *Y. aloifolia* and *Y. filamentosa*, respectively (Fig. 2, Supplementary Fig. S2, Supplementary Tables 4 and 5). Transcription factors (TFs) were notably scarce in our proteomic dataset with only 256 out of 5,636 annotated TFs observed in *Y. aloifolia* and 349 out of 5,521 annotated TFs observed in *Y. filamentosa*.

Untargeted metabolomic analysis identified 72 metabolites in *Y. aloifolia* and 81 metabolites in *Y. filamentosa* of which 48 were found in both species (Supplementary Table 6). In *Y. aloifolia* 21/72 metabolites showed time-structured abundance over the day-night period in at least one treatment compared to 10/81 in *Y. filamentosa* (Supplementary Table 6). Of the metabolites identified in both species, just 5/48 metabolites cycled in at least one treatment in both species (Supplementary Table 6).

### The CAM pathway

Of CAM associated proteins (Fig. 3a), relatively few exhibited cycling abundance across time. While no copies of β-CA cycled in *Y. aloifolia,* two copies cycled in *Y. filamentosa*, one of which had no clear ortholog in *Y. aloifolia.* (Fig. 3b, Supplementary Fig S3). Notably, *Y. aloifolia* exhibits cycling mRNA expression of two distinct forms of PEPC (Heyduk et al. 2022) commonly referred to as plant-type PEPC (PTPC), and bacterial-type PEPC (BTPC; Supplementary Fig. S3). Two copies of PTPC cycled in each *Y. aloifolia* and *Y. filamentosa* with one cycling in both. Curiously, the copy in *Y. aloifolia* equal-contribonding to the most highly expressed mRNA transcript – typically associated with CAM – did not cycle in protein abundance, while its ortholog in *Y. filamentosa* did. In the CAM pathway, only BTPC protein abundance both cycled and appeared to track mRNA expression under drought and well-watered conditions in *Y. aloifolia* where two copies showed similar cycling abundance (Fig 3b, Supplementary Fig. S3). BTPC abundance did not cycle in *Y. filamentosa*. PPCK, responsible for phosphorylating PEPC to enhance its activity, was notably absent from the proteomic data in both species, potentially due to its low abundance (Hartwell et al. 1999). We found 2 copies of cytoplasmic NAD-MDH that cycled in abundance in *Y. filamentosa* but none in *Y. aloifolia* (Fig 3b, Supplementary Fig. S3). In the light reactions, CAM associated copies of NADP-ME and PPDK cycled in protein abundance in *Y. aloifolia* under well-watered conditions, though peak expression was shifted 4 to 8 hours later in both relative to mRNA expression (Fig. 3b). The same copy of NADP-ME also cycled in *Y. filamentosa* under drought with peak abundance just before dusk, despite the fact that its transcript did not cycle. A second paralog of NADP-ME cycled in *Y. aloifolia* with peak protein abundance during the dark period despite its hypothesized involvement in the daytime decarboxylation reactions in CAM (Supplementary Fig. S3).

In contrast to protein abundance, the majority of CAM associated mRNA transcripts exhibited cycling expression in *Y. aloifolia* (33/38) relative to *Y. filamentosa* (18/36; Supplementary Table 6). As observed in previous studies (Heyduk et al., 2019), *Y. aloifolia* showed coordinated cycling of canonical CAM genes, including PEPC, PPCK, NADP-ME, and PPDK, equal-contribonding to their expected roles in nocturnal carboxylation and daytime decarboxylation. While expression phase differed among some paralogs and treatments (Fig. 3b, Supplementary Fig. S3), overall patterns support a reliance on the NADP-ME/PPDK decarboxylation route. In *Y. filamentosa*, CAM-related transcripts were generally expressed at lower levels and cycled less frequently, consistent with its reliance on C_3_ photosynthesis.

Malate was the only core CAM metabolite present in our metabolomic dataset. As expected, malate cycled strongly in *Y. aloifolia* relative to *Y. filamentosa*, accumulating throughout the night and decreasing in abundance during the day (Fig. 3c). Despite some similarity between species in expression of CAM genes such as *PPCK* and at least one copy of *PTPC*, no nighttime accumulation of malate was observed in *Y. filamentosa*. This result was consistent with measurements of nocturnal stomatal conductance in *Y. aloifolia* relative to *Y. filamentosa* (Supplementary Fig. S1). In *Y. aloifolia*, but not *Y. filamentosa*, phosphate abundance also cycled strongly and tracked malate abundance, increasing at night and decreasing during the day (Fig. 3c). Cycling of phosphate in *Y. aloifolia* likely correlates with its release during the carboxylation of PEP to OAA at night, its essential role in activation of PEPC, and high demand during the day for regenerating compounds such as ATP and NADP.

### Citrate accumulation

Citrate exhibited similar fluctuation in abundance to malate in *Y. aloifolia*, as previously reported in several other CAM taxa (Franco et al. 1992; Freschi et al. 2010; Borland and Griffiths 1989) leading us to investigate genes associated with citrate metabolism in both species (Fig. 4). In plants, citrate is typically produced by citrate synthase (CS) in the peroxisome as part of the glyoxylate cycle and in the mitochondria, as part of the tricarboxylic acid cycle (TCA) cycle. In the mitochondria, citric acid is subsequently converted to isocitrate and isocitrate to 2-oxoglutarate (2-OG) by aconitase (ACO) and NAD-dependent isocitrate dehydrogenase (NAD-IDH), respectively (Fig. 4a). Abundance of citrate in *Y. aloifolia* increased throughout the night and decreased during the day, compared to *Y. filamentosa* where citrate levels were relatively stable between day and night (Fig. 4c).

**Figure 4:**
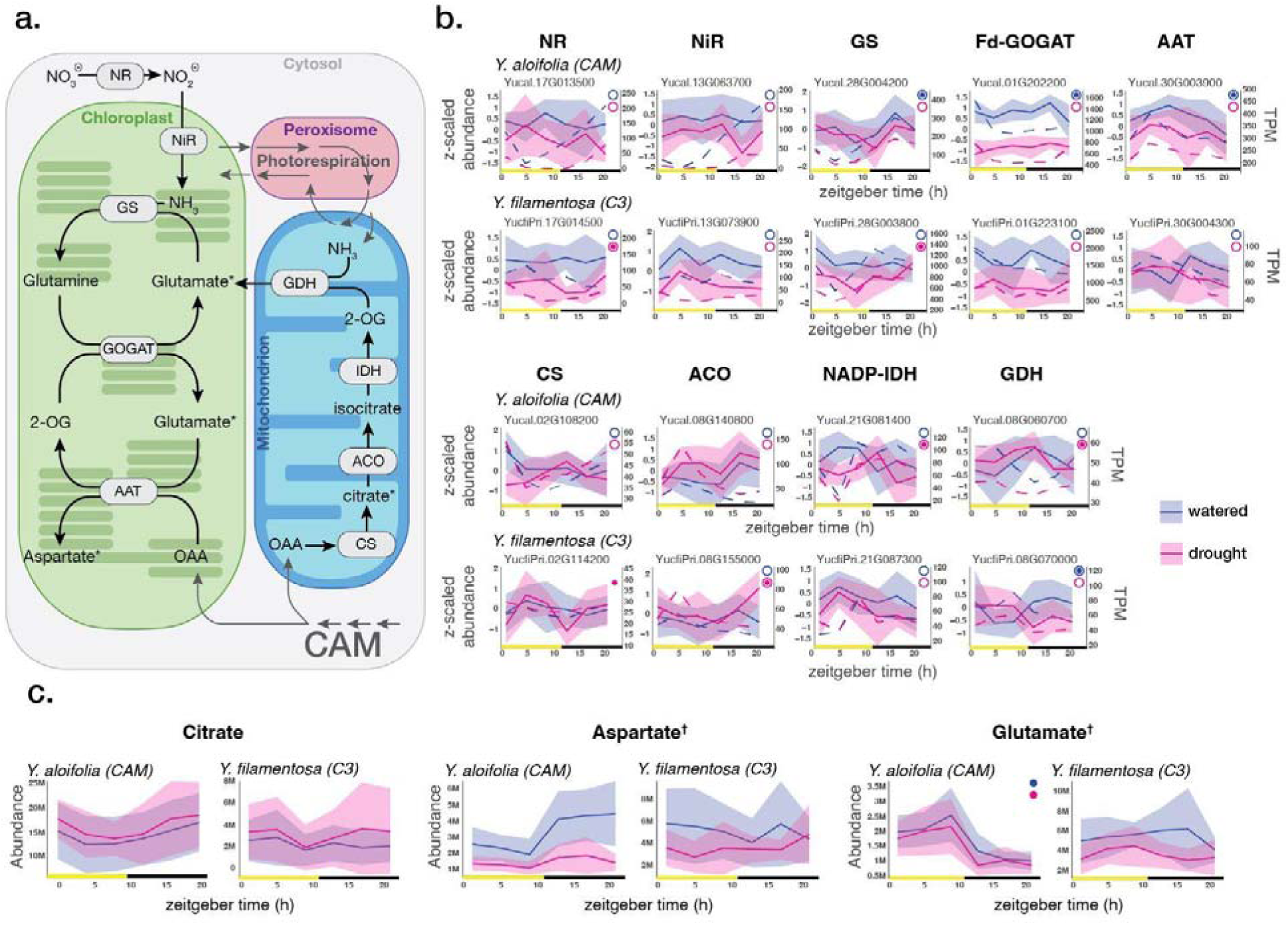
Comparison of nitrogen metabolism in *Y. aloifolia* and *Y. filamentosa.* **(a.)** Flow-chart of core reactions of the nitrogen metabolism pathway and its interactions with CAM, photorespiration, and the citric acid cycle. Conversion of citrate to 2-OG may occur in the cytosol (not shown) or mitochondrion (as shown). **(b.)** Plots of z-scaled mean protein abundance (solid line, SD represented by shaded area) with mean expression in TPM overlaid (dashed line) for comparison. Filled and unfilled circles indicate significantly (*p* < 0.05) time-structured protein abundance and gene expression, respectively. **(c.)** Plots of abundance over time for glutamate, aspartate, and citrate (indicated in flow-chart by an ‘*’) shaded areas represent the standard deviation. Filled circles indicate significantly (*p* < 0.05) time-structured metabolite abundance and daggers (†) indicate metabolites that are significantly (*p* < 0.05) different between species. Abbreviations: 2-OG, 2-oxoglutarate; AAT, aspartate aminotransferase; ACO, aconitase; CS, citrate synthase; GDH, glutamate dehydrogenase; GOGAT, glutamine oxoglutarate aminotransferase; GS, glutamine synthetase; IDH, isocitrate dehydrogenase; NADP-IDH, NADP-dependent isocitrate dehydrogenase; NiR, nitrite reductase; NR, nitrate reductase; OAA, oxaloacetate.

As in core CAM pathway genes, relatively few proteins associated with citrate production cycled in abundance in either *Y. aloifolia* or *Y. filamentosa* and even fewer were associated with transcripts that also cycled in expression. A single copy of mitochondrial CS protein cycled under drought in *Y. aloifolia*, but did not equal-contribond to the most highly expressed cycling transcript in that species (Fig 4b, Supplementary Fig. S4). One mitochondrial and one peroxisomal copy of CS protein appeared to cycle in *Y. filamentosa* under drought conditions. However both exhibited shifts in their abundance profiles relative to transcript expression, particularly under drought-stress. ACO followed a similar trend, in that the most highly expressed cycling transcripts were not associated with cycling protein abundance, though one gene in *Y. aloifolia* and two in *Y. filamentosa* exhibited cycling protein abundance under at least one treatment (Fig 4b, Supplementary Fig. S4, Supplementary Table 6). The most highly expressed copy of NADP-IDH cycled under drought in *Y. aloifolia* but not in *Y. filamentosa*.

In contrast to our proteomic data, 35 out of 40 TCA cycle genes cycled in transcript expression in *Y. aloifolia* and exhibited a diverse array of expression patterns, while the relatively few transcripts that cycled in *Y. filamentosa* showed highly similar expression patterns in our MaSigPro analysis (cluster 1, Supplementary Fig. S4, Supplementary Table 6). Notably, in *Y. aloifolia*, one copy of *CS*, two copies of *ACO*, one copy of *NAD-IDH*, and one copy of *NADP-IDH* all had similar cycling expression that tracked temporal patterns of citrate abundance with peak expression during the dark period (Fig. 4b, Supplementary Fig. S4). The most highly expressed copy of mitochondrial *CS* cycled in *Y. aloifolia* but not in *Y. filamentosa*, exhibiting peak expression at dawn. Two copies of *ACO* exhibited strong cycling expression in *Y. aloifolia*, with expression decreasing during the morning before increasing and remaining high throughout the night (Fig. 4b, Supplementary Fig. S4). Expression of *ACO* in *Y. filamentosa* did not have a clear temporal pattern and was 3-4 times lower than in *Y. aloifolia*. Differences in expression were further observed in both NAD-dependent (mitochondrial) and NADP-dependent (cytosolic) copies of *IDH*. *NAD-IDH* expression was relatively stable over time in both species and treatments but roughly twice as high in *Y. aloifolia* (Supplementary Fig. S4). Cytosolic *NADP-IDH* exhibited peak expression in both species near dusk but remained high throughout the night in *Y. aloifolia*, strongly resembling the expression of *ACO* (Fig. 4b).

### Nitrogen metabolism

Outside of the CAM pathway, our multi-omics analysis allowed us to explore differences in protein abundance and gene expression associated with nitrogen metabolism. Briefly, nitrogen metabolism begins with the reduction of nitrate to ammonia via nitrate reductase (NR) and nitrite reductase (NiR), followed by its incorporation into glutamine and glutamate through the glutamine synthetase/glutamate synthase (GS/GOGAT) cycle in the chloroplast (Fig. 4a). Aspartate aminotransferase (AAT) links nitrogen assimilation and carbon metabolism by transferring amino groups between glutamate and oxaloacetate to yield aspartate and 2-oxoglutarate (2-OG), while glutamate dehydrogenase (GDH) catalyzes the reversible conversion of glutamate, ammonia, and 2-OG in the mitochondrion, providing another connection between nitrogen metabolism and the TCA cycle (Fig. 4a).

Multiple proteins were found to cycle in abundance in *Y. aloifolia* and to a lesser extent in Y. filamentosa. Two notable exceptions were found in NR and NiR, which catalyze the initial reduction of nitrate to ammonia. Neither NR nor NiR cycled in abundance in *Y. aloifolia*, though one copy of NR did cycle in *Y. filamentosa*, peaking in abundance at midday (Fig. 4b). In the chloroplast, multiple copies of GS exhibited cycling protein abundance in both species, while ferredoxin-dependent glutamate synthase (Fd-GOGAT) and AAT each had multiple paralogs that cycled in *Y. aloifolia* alone (Fig. 4b, Supplementary Fig. S5, Supplementary Table 6). A single copy of GDH cycled in protein abundance in both species but showed inverse patterns of protein abundance, peaking at dawn in *Y. aloifolia* and just before dusk in *Y. filamentosa* (Fig. 4b). One copy of GS exhibited remarkably similar cycling abundance between species, peaking during the dark period in *Y. aloifolia* in well-watered and drought conditions as well as *Y. filamentosa* under drought (Fig. 4b). Another copy of GS cycled in abundance in *Y. aloifolia* under well-watered conditions and two copies cycled in *Y. filamentosa* under well-watered and drought conditions (Supplementary Fig. S5, Supplementary Table 6). Both copies of Fd-GOGAT exhibited cycling abundance in *Y. aloifolia* (Fig. 4b, Supplementary Fig. S5, Supplementary Table 6) with higher abundance during the dark period, but not in *Y. filamentosa*. Similarly, 4 out of 5 AAT proteins cycled in *Y. aloifolia* with peak expression near dusk (Fig. 4b, Supplementary Fig. S4, Supplementary Table 6). None of the copies of AAT present in *Y. filamentosa* displayed time structured protein abundance.

Several nitrogen metabolism genes showed clear differences in transcript expression between species and treatments, even in cases where their equal-contribonding proteins did not (Fig. 4b, Supplementary Fig. S4, Supplementary Table 6). For example, *NR* and *NiR*, cycled in Y. aloifolia, peaking near dawn under both treatments, whereas in Y. filamentosa cycling occurred only under drought (Fig. 4b). *GDH* also showed distinct temporal expression, peaking near dawn in *Y. aloifolia* and around midday in *Y. filamentosa* in both treatments with peak gene expression shifted 8 hours earlier than equal-contribonding protein abundance in *Y. aloifolia* and similarly under well-watered conditions in *Y. filamentosa* (Fig. 4b). In contrast, other genes in the nitrogen metabolism pathway, including *GS*, *Fd-GOGAT*, and *AAT*, showed similar cycling expression patterns across species. In addition, two copies of *AAT* and *Fd-GOGAT* cycled in *Y. filamentosa* transcripts despite none of the proteins cycling in abundance (Fig. 4b, Supplementary Fig. S5, Supplementary Table S6). Expression levels of *GS* and *Fd-GOGAT* were generally lower in *Y. aloifolia*, especially under well-watered conditions, while *AAT* expression was nearly four times higher in *Y. aloifolia* across both treatments (Fig. 4b).

Two primary products of nitrogen metabolism, aspartate and glutamate, were identified by metabolomic analysis (Fig 4c). The production of aspartate and glutamate by AAT and Fd-GOGAT, respectively, plays a central role in primary nitrogen metabolism and uses OAA, a primary product of CAM dark reactions and TCA cycle intermediate, as a substrate. In Y. aloifolia, aspartate and glutamate exhibited inverse abundance patterns, with aspartate peaking at night and glutamate peaking during the day (Fig. 4c). Despite this striking difference in metabolite abundance, expression patterns of *AAT* and *Fd-GOGAT* were broadly similar between CAM and C_3_ species, though overall expression of *AAT* was considerably higher in *Y. aloifolia*. In addition, the most highly expressed copies of AAT and Fd-GOGAT cycled in protein abundance under well watered conditions in *Y. aloifolia* but not in *Y. filamentosa*, peaking at the end of the day (Fig. 4b). Conversely, *GDH*, which potentially influences glutamate availability by its interconversion with 2-oxoglutarate and ammonia in the mitochondrion, displayed distinct cycling patterns of expression and abundance between the two species.

### Photorespiration

Very few genes in the photorespiratory pathway (Fig. 5a) displayed cycling protein abundance in either species of *Yucca*. Out of 8 core genes involved in photorespiration only mitochondrial serine hydroxymethyl transferase (SHMT) and glycine decarboxylase complex - subunit H (GDC-H) cycled in protein abundance in both species (Fig. 5b, Supplementary Fig. S6), under well-watered conditions in *Y. aloifolia* and drought in *Y. filamentosa*. SHMT and the GDC enzyme complex facilitate the conversion of glycine to serine in the mitochondria as part of the photorespiratory pathway (Fig. 5a). Specifically, as a component of the GDC enzyme complex, GDC-H plays an essential role in the decarboxylation of glycine and subsequent release of photorespiratory CO_2_ and ammonia (Timm et al. 2012).

**Figure 5:**
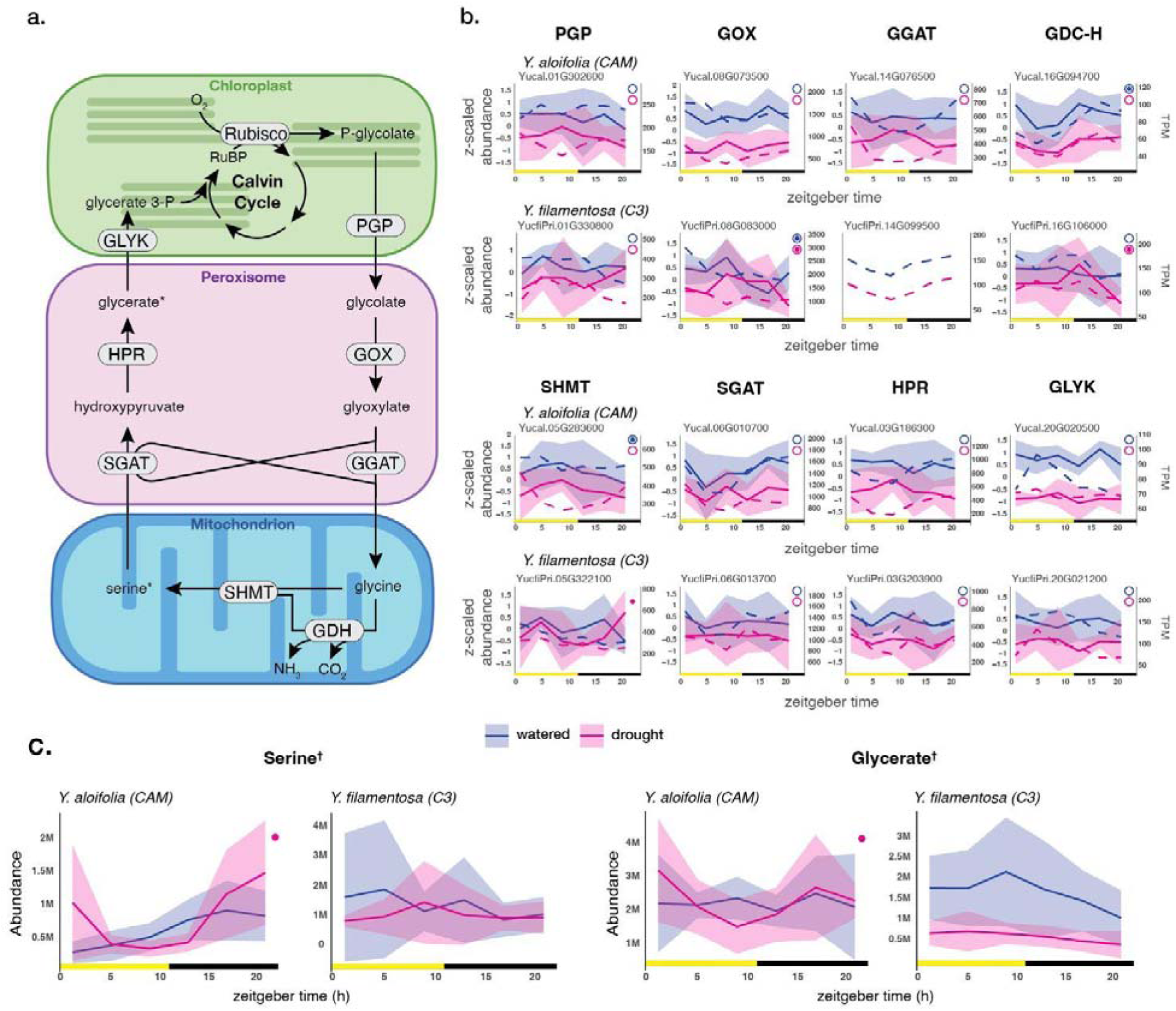
Comparison of photorespiration in *Y. aloifolia* and *Y. filamentosa.* **(a.)** Flow-chart of core reactions of the photorespiratory pathway. **(b.)** Plots of z-scaled mean protein abundance (solid line, SD represented by shaded area) with mean expression in TPM overlaid (dashed line) for comparison. Filled and unfilled circles indicate significantly (*p* < 0.05) time-structured protein abundance and gene expression, respectively. **(c.)** Plots of abundance over time for serine and glycerate (indicated in flow-chart by an ‘*’) shaded areas represent the standard deviation. Filled circles indicate significantly (*p* < 0.05) time-structured metabolite abundance and daggers (†) indicate metabolites that are significantly (*p* < 0.05) different between species. Abbreviations: GDH/GDC-H, glycine decarboxylase complex - subunit H; GGAT, glutamate:glyoxylate aminotransferase; GLYK, glycerate kinase; GOX, glycolate oxidase; HPR, hydroxypyruvate reductase; PGP, phosphoglycolate phosphatase; Rubisco, ribulose bisphosphate carboxylase; SGAT, serine:glyoxylate aminotransferase; SHMT, serine hydroxymethyl transferase.

Overall transcript expression of genes in the photorespiratory pathway was similar between *Y. aloifolia* and *Y. filamentosa* with many genes peaking near dawn or during the dark period (Fig. 5b). A notable exception was phosphoglycolate phosphatase (*PGP*), the enzyme that catalyzes the first step of the photorespiratory pathway, converting 2-phosphoglycolate to glycolate in the chloroplast. Two copies of *PGP*, were expressed in *Y. filamentosa*, and both had elevated expression throughout the daylight hours (Fig 5b., Supplementary Fig. S6), consistent with elevated photorespiratory stress during the day. Conversely, *Y. aloifolia*exhibits treatment specific expression of a single copy of *PGP*. Under well-watered conditions, *PGP* peaks at midday and again during the night. Under drought, PGP steadily increases at night, peaking at dawn (Fig. 5b). Most cycling photorespiratory genes clustered consistently within species, with peak expression at the end of the night in *Y. aloifolia* (15/24 transcripts, including 6/8 highly expressed copies) and *Y. filamentosa* (10/20 transcripts). An additional cluster (5/20 transcripts) with peak expression at midday was observed only in *Y. filamentosa*. Serine hydroxymethyltransferase (SHMT) showed unusually diverse expression, with most paralogs in both species falling outside the dominant photorespiratory clusters (Supplementary Fig. S6, Supplementary Table 6).

*Yucca aloifolia* and *Y. filamentosa* exhibited distinct metabolite abundance for serine and glycerate, produced as intermediates in the photorespiratory pathway (Fig. 5a). In Y. aloifolia, both serine and glycerate cycled in abundance during drought showing elevated abundance during the night. In contrast, *Y. filamentosa* showed steady abundance of serine and glycerate under drought and well watered conditions (Fig. 5c). Glycolate, produced by *PGP* in the first step of the photorespiratory cycle, was also detected in *Y. aloifolia* but not in *Y. filamentosa* and did not cycle.

## Discussion

Through the use of integrated -omics data, this study takes advantage of two closely-related (McKain et al. 2016) but photosynthetically distinct species of *Yucca* to understand the regulation of CAM photosynthesis at the cellular level. As expected based on prior work (Heyduk et al. 2019), *Y. aloifolia* has nocturnal CO_2_ assimilation (Supplementary Fig. S1) (though, notably, daytime CO_2_ assimilation is not zero) and a significant accumulation of malate over the course of the night (Fig. 3c). In contrast, *Y. filamentosa* does not take up CO_2_ at night (Supplementary Fig. S1), nor does it have a significant accumulation of malate under either treatment condition (Fig. 3c). These species thus provide an ideal framework for comparing transcription, metabolite profiles, and protein abundance across divergent photosynthetic pathways. Our results highlight core differences in nitrogen metabolism and photorespiration between the CAM and C_3_ species, and also suggest relatively low dynamic changes to protein abundance over the course of the diel cycle in contrast to transcript abundance.

### Protein abundance shows reduced temporal expression relative to mRNA

Relatively few proteins cycled in abundance and protein abundance rarely cycled along with their associated transcripts, or in cases where they did, they frequently exhibited shifts in peak abundance relative to mRNA expression (Fig. 3). The lack of correlation between proteins and their transcripts was particularly apparent in the core CAM pathway, to the extent that it might be difficult to distinguish between CAM and C_3_ on the basis of protein abundance alone (Fig 3; Supplementary Fig. S3). In any case, our results align with previous studies that found relatively low correlation between mRNA expression and protein abundance in other plants (de Sousa Abreu et al. 2009; Kundrátová et al. 2021; Baerenfaller et al. 2012; Guan et al. 2024; Schiller et al. 2025). The lack of correlation between transcript expression and protein abundance is potentially driven by multiple factors. First, far fewer gene products were represented in our protein abundance dataset relative to our transcript data. Many genes with low expression (below 100 TPM average) did not have associated protein data. As a result, it is possible that some correlations were not detected. For example, proteins that are commonly present in low abundances including transcription factors (e.g. PRR1, CCA1/LHY) and protein kinases (e.g. PPCK), were absent from our proteomic dataset regardless of their expression at the transcript level. Many of these proteins are key regulators of metabolism that are associated with transcripts that cycle in expression throughout the day and are similarly expected to exhibit strong cycling abundance themselves (Hartwell et al. 1999). Second, there is ample evidence that shifts in protein abundance are driven by factors other than mRNA expression including post-transcriptional regulation, translation, and degradation of protein products (Vogel and Marcotte 2012; Wu et al. 2024). Finally, it is important to keep in mind that metabolic rates are not solely regulated by protein abundance. Processes such as protein-protein interaction (Zhang et al. 2018; Rugen et al. 2021), allosteric regulation by metabolites (Hartman et al. 2023), and post-translational modification (PTM) have significant effects on primary metabolism (Balparda et al. 2023). For instance, several proteins in the CAM pathway, including PEPC (Giglioli-Guivarc’h et al. 1996), MDH (Carr et al. 1999), and PPDK (Chen et al. 2014), are subject to post-translational interactions and modifications that affect their activity. Likewise, multiple enzymes involved in nitrogen metabolism (Seabra et al. 2013; Costa-Broseta et al. 2020; Kaiser and Huber 2001) and photorespiration (Palmieri et al. 2010; Jossier et al. 2019; Liu et al. 2020) are known to be regulated by PTMs.

### Modification of primary metabolism in Y. aloifolia

The evolution of CAM involves temporal shifts in gene expression and primary metabolite accumulation. Broad modifications to core processes associated with carbon fixation likely necessitate simultaneous adjustments across other pathways that share metabolic substrates and overlap in regulation (Fig. 1). In particular, PEPC activity is strongly associated with N-metabolism in C_3_ plants as it is responsible for replenishing carbon skeletons diverted from the TCA cycle for amino acid synthesis (O’Leary and Plaxton 2015). Our investigation into downstream effects of CAM in *Yucca* reveals significant differences in N-metabolism between *Y. aloifolia* and *Y. filamentosa* (Fig. 4, Supplementary Fig. S4), in agreement with previous studies highlighting links between CAM photosynthesis and nitrogen assimilation in various CAM and C_3_+CAM lineages (Gonçalves and Mercier 2021; Santos and Salema 1991; Freschi et al. 2010; Pereira et al. 2018). Under well-watered conditions, we observed changes in gene expression related to nitrate reduction and amino acid synthesis (Fig 4b, Supplementary Fig. S4). Additionally, there were clear differences in accumulation of glutamate and aspartate, immediate downstream products of primary nitrogen metabolism (Fig. 4c). Accumulation of glutamate during the light period in *Y. aloifolia* was associated with a shift in expression of GDH from midday to early morning. Conversely, the sharp increase in aspartate abundance coupled with the drop in glutamate abundance overnight was not accompanied by any changes in expression of AAT, which can interconvert OAA and glutamate with 2-OG and aspartate. Thus, increased nocturnal production of OAA may be sufficient to explain nocturnal synthesis of aspartate via AAT without alterations to transcript expression or protein abundance. Prior research has shown that, along with malate, aspartate is a major transporter of assimilated carbon into the bundle sheath cells of C_4_ plants (Meister et al. 1996; Schlüter et al. 2019). Additionally, aspartate transport in and out of the vacuole has been demonstrated in C_3_ plants (Snowden et al. 2015). Accordingly, it is plausible that at least some CAM plants including *Y. aloifolia* may use aspartate as an alternative form of carbon storage during the night, supplementing or buffering malate-based carbon pools during CAM.

Alongside well-established differences in expression in genes associated with the central CAM carboxylation and decarboxylation pathways, we observed differences in accumulation of organic acids other than malate, such as citrate, malonic acid, and maleic acid in the CAM species *Y. aloifolia* relative to the C_3_ species *Y. filamentosa* (Fig. 4c, Supplementary Table 5). Nocturnal accumulation of citrate has been documented in multiple CAM species (Franco et al. 1992; Kornas et al. 2009; Freschi et al. 2010; Borland and Griffiths 1989; Lüttge 1988, 1990). However, its role in CAM is less clear than that of malate, as fixation of CO_2_ into citrate does not result in a net carbon gain (Lüttge 1988). Several theories have been put forth regarding putative benefits of citrate accumulation in CAM plants. In the CAM pathway, the production of malate by MDH is dependent on NADH, as opposed to citrate synthesis which results in a net gain of NADH. As such, nocturnal citrate production may provide additional reducing power for reduction of malate while also serving as a secondary storage mechanism for excess CO_2_ as internal stores of NADH are exhausted (Trípodi and Podestá 2003). Additionally, in *Yucca*, we observed a difference in peak gene expression of NR and NiR from midday in *Y. filamentosa* to the end of the night in *Y. aloifolia*. This may suggest that the additional reducing power generated by citrate synthesis is driving reduction of nitrate during the dark period in *Y. aloifolia,* as opposed to C_3_ plants which primarily generate reducing power for N-assimilation via photosynthesis during the day (Foyer et al. 2001). It is also possible that accumulation of citrate is merely an atavistic trait retained in CAM plants from their C_3_ ancestors (Bräutigam et al. 2017). However, this scenario seems unlikely as we do not observe any nocturnal accumulation of citrate in *Y. filamentosa*.

In C_3_ species, the decarboxylation of citrate during the day is thought to supply the carbon skeletons for amino acid synthesis (Scheible et al. 2000). Citrate’s role in linking carbon and nitrogen metabolism in C_3_ plants may offer an explanation for the inverse cycling of aspartate and glutamate observed in *Y. aloifolia* throughout the day and night (Fig 4c). In C_3_ plants, 2-OG is supplied for nitrogen assimilation from the decarboxylation of citrate in the mitochondria or the cytosol (Szal and Podgórska 2012). Additionally, there is evidence that 2-OG used for daytime nitrogen assimilation in C_3_ plants originates, at least in part, from citrate accumulated during the night (Tcherkez et al. 2009). As 2-OG is the direct substrate for glutamate synthesis by GDH and GOGAT, the high accumulation of glutamate during the day in *Y. aloifolia* relative to *Y. filamentosa* could be driven by the increase in citrate production during the dark period and its subsequent degradation during the day (Fig. 4).

In both CAM and C_3_ plants, there is a clear association between citrate production and various types of abiotic stress (Tahjib-Ul-Arif et al. 2021; Lüttge 2006). Citrate:malate ratios are known to increase in CAM plants in response to high light stress in *Tillandsia* (Borland and Griffiths 1989) and high light stress and drought in *Clusia* (Franco et al. 1992). Citrate production is more energetically favorable than that of malate, as the conversion of hexose to citrate results in a net gain of ATP and NAD(P)H relative to fixation of CO_2_ into malate (Lüttge 1988). In addition, the conversion of citric acid to OAA releases two molecules of CO_2_ whereas the conversion of malate to pyruvate during the light reactions of CAM releases only a single molecule of CO_2_. As such, production of citrate may represent a way to recycle respiratory CO_2_ during periods of drought where stomatal conductance is severely reduced during the day and night, limiting access to atmospheric CO_2_ (Lüttge 1988). Likewise, under high light, accumulation of citrate at night and the subsequent release of CO_2_ upon its breakdown during the day is thought to minimize photorespiration and photoinhibition by concentrating CO_2_ around Rubisco as supplies of malate are exhausted (Miszalski et al. 2013; Lüttge 1990). Finally, it has been suggested in *Clusia* that accumulating citrate at night could increase the buffering capacity of the vacuole, leading to an increase in storage capacity for CO_2_ (Franco et al. 1992). Our observations in *Y. aloifolia* were consistent with those in other species that accumulate citrate with the ratio of citrate to malate increasing under stress (Franco et al. 1992; Lüttge 1990). This may be due to citrate’s role in minimizing oxidative stress or alternatively may reflect the lower energetic requirements of citrate production relative to malate (Lüttge 1988). Regardless, our findings highlight a potentially important link between citrate metabolism and CAM in *Yucca*.

Analysis of gene expression and protein abundance alongside fluctuations in metabolite abundance demonstrate a clear effect on overlapping pathways of nitrogen assimilation and to a lesser extent, photorespiration. The CAM metabolite OAA seems to play a key role in driving shifts in primary metabolism and may act as a link between regulation of carbon and nitrogen metabolism (Chen et al. 2022). Multiple metabolites that exhibit day-night fluctuations in abundance in *Y. aloifolia*, including those in the nitrogen cycle, rely on OAA as a substrate (Fig.1). The cycling abundance of these primary metabolites is notable as OAA is produced in great abundance by carboxylation of PEP by the enzyme PEPC in CAM plants. The subsequent reduction of OAA to malate by MDH is dependent on the presence of NADH (Trípodi and Podestá 2003). Under certain conditions, it is possible that OAA synthesis by PEPC outstrips the ability of MDH to convert OAA to malate. This could potentially result in OAA being funneled into other processes that either generate reducing power, such as citrate synthesis, or at the very least do not require NADH, such as aspartate synthesis. As such, cycling of multiple metabolites outside of the central CAM pathway may initially be driven by greater flux through OAA rather than extensive regulatory rewiring outside of the core CAM pathway.

### Photorespiratory pathway largely conserved between CAM and C_3_ plants

The mitigation of photorespiration is thought to play a major role in CAM evolution; by using PEPC as the initial carboxylating enzyme to concentrate carbon around Rubisco, CAM plants are thought to reduce photorespiration by preventing Rubisco’s interaction with O_2_. It is therefore assumed that photorespiration decreases alongside the evolution of carbon concentrating mechanisms like CAM and C_4_. However, the photorespiratory pathway itself has only rarely been the direct focus of CAM research, and studies have shown conflicting results. In a strong CAM species, internal concentrations of CO_2_:O_2_ were high enough during the day to largely suppress photorespiration, while a facultative CAM species demonstrated high levels of photorespiration even when CAM was induced (Spalding et al. 1979; Duarte and Lüttge 2007). Our results show that while there are subtle differences in gene expression and protein abundance related to photorespiratory pathways between CAM and C_3_ species of *Yucca*, both remain largely conserved between species in drought and well-watered conditions. A notable exception to this is found in gene expression, though not necessarily protein abundance, of PGP, which has daytime expression in *Y. filamentosa* but is highly expressed at night in *Y. aloifolia* under both treatments. The difference is particularly pronounced under drought where expression of PGP is nearly inverse between species. Conserved expression of multiple photorespiratory genes between *Y. aloifolia* and *Y. filamentosa* combined with altered expression of PGP in *Y. aloifolia* may suggest that different portions of the photorespiratory pathway are under distinct regulatory control. Furthermore, it is possible that differences in expression of PGP, particularly under drought, could be linked to reduced photorespiration in *Y. aloifolia*.

Despite broad similarities in gene expression and protein abundance between species and treatments, we observed nocturnal accumulation of glycerate and serine, two products of the photorespiratory pathway, exclusively under drought in *Y. aloifolia* (Fig. 5c). In addition, we did not observe strong accumulation of serine or glycerate during the day in either species under either treatment in contrast to what has been observed in other C_3_ plants (Sato et al. 2008; Timm et al. 2013). Serine is an important substrate for 1C metabolism in plants where it serves as a major donor of one-carbon units for the biosynthesis of nucleotides, amino acids, and other essential metabolites. Serine is primarily produced via photorespiration in C_3_ plants (Ros et al. 2014). However, the nocturnal accumulation of serine in *Y. aloifolia* is unlikely to be driven by photorespiratory flux, as photorespiration is a light-dependent process (Foyer et al. 2009). Alternative pathways of serine biosynthesis exist in plants commonly referred to as the “phosphorylated serine pathway” and the “non-phosphorylated (glycerate) serine pathway” (Igamberdiev and Kleczkowski 2018). Based on the time structured nature of both glycerate and serine under drought, along with the absence of phosphorylated intermediates in our metabolomic data, nocturnal accumulation of serine and glycerate in *Y. aloifolia* could be driven by the poorly characterized, non-phosphorylated serine pathway of serine biosynthesis. This pathway is thought to be an important source of serine in C_4_ plants where photorespiration is greatly reduced (Kleczkowski and Givan 1988; Randall et al. 1971). Many obligate CAM plants such as *Y. aloifolia* open their stomata during the early morning and late evening (Males and Griffiths 2017), providing an opportunity for normal, C_3_ levels of photorespiration to occur at the beginning and end of day. However, drought further reduces stomatal conductance (Herrera et al. 2000), restricting gas exchange entirely to the dark period in *Y. aloifolia* (Supplementary Fig. S1) and essentially eliminating fixation of O_2_ by Rubisco. Thus, it is likely that nocturnal accumulation of serine and glycerate under drought in *Y. aloifolia* does not reflect flux through the photorespiratory pathway, but production of serine by an alternative pathway. It is also notable that serine has been shown to be a potentially powerful transcriptional regulator of multiple genes in the photorespiratory pathway in C_3_ plants (Timm et al. 2013). Specifically, external application of serine has been shown to deregulate the expression of GDC, SHMT, and GLYK, resulting in significantly higher expression during the night than the day in Arabidopsis. Expression of PGP was also altered but remained higher during the day. Additionally, the deregulation of expression of photorespiratory genes resulted in substantial discordance between expression of transcripts and abundance of their equal-contribonding protein products (Timm et al. 2013). This pattern is similar to what we observe in *Y. aloifolia* and *Y. filamentosa*, potentially indicating a link between non-photorespiratory serine production and alterations to gene expression and protein abundance in the photorespiratory pathway in *Yucca*. However, the extent to which this might be tied to the evolution of CAM in *Yucca* remains unclear.

It is also of interest that when *Y. filamentosa* experiences drought, the levels of aspartate and glutamate have similar abundance patterns as those in *Y. aloifolia* (Fig. 3c). However, the similarity between C_3_ and CAM species in metabolite abundance under drought does not extend to photorespiratory metabolites found in our data. This may suggest that the evolution of CAM was not primarily driven by shifts in the photorespiratory pathway, as drought induces "CAM-like" behavior in nitrogen metabolism but not photorespiration. Instead, CAM evolution may only initially require changes to expression of a handful of genes, which primarily enhance metabolic flux through existing C_3_ pathways rather than requiring complex, global regulatory shifts across multiple pathways. Additional, detailed comparisons of molecular time series and functional analysis of photorespiratory genes in other related CAM and C_3_ species are needed to fully understand the dynamics we observe in *Y. aloifolia* and *Y. filamentosa*.

### Considerations for future studies in CAM evolution

Our study shows the importance of comparative multi-omic analysis between closely related C₃ and CAM taxa and underscores the need to expand this work across diverse lineages, including aquatic and physiologically intermediate C₃ + CAM species. While transcriptomic data remain valuable, the limited concordance between transcript, protein, and metabolite abundance in *Yucca* emphasizes that relying on transcriptomic data alone provides a limited and potentially biased view of underlying molecular processes. Future studies should strive to employ a broader range of molecular data to more fully resolve the regulatory complexity underlying CAM. In particular, combining analyses of protein and metabolite abundance with phospho-proteomics and advanced genomic assays such as ATAC-seq and ChIP-seq will provide powerful means of probing the diverse regulatory mechanisms that govern CAM expression and evolution. Furthermore, comparative genomics alone cannot reliably infer gene function. Uncovering the role of specific genes will ultimately require functional validation through experimental transformation in CAM and related C₃ lineages. By identifying promising gene targets beyond the core CAM pathway, our work lays the foundation for a transition from broad comparative analyses to such targeted and experimental approaches capable of uncovering the mechanisms driving the convergent evolution of CAM across vascular plants.

## Supplementary Data

*Supplementary Figure 1:* Stomatal conductance varies by treatment and condition in *Yucca*.

*Supplementary Figure 2:* Detailed comparison of cycling transcripts and proteins in *Y. aloifolia* and *Y. filamentosa*.

*Supplementary Figure 3:* Detailed mRNA expression and protein abundance data for all paralogs of genes in the CAM pathway.

*Supplementary Figure 4:* Detailed mRNA expression and protein abundance data for all paralogs of genes associated with citrate metabolism.

*Supplementary Figure 5:* Detailed mRNA expression and protein abundance data for all paralogs of genes associated with nitrogen metabolism.

*Supplementary Figure 6:* Detailed mRNA expression and protein abundance data for all paralogs of genes associated with photorespiration.

*Supplementary Table 1:* Summary of maSigPro results and peak mean expression for *Y. aloifolia*.

*Supplementary Table 2:* Summary of maSigPro results and peak mean expression for *Y. filamentosa*.

*Supplementary Table 3:* Results of linear modeling on protein data for *Y. aloifolia*.

*Supplementary Table 4:* Results of linear modeling on protein data for *Y. filamentosa*.

*Supplementary Table 5:* Summary of metabolomics data for *Y. aloifolia* and *Y. filamentosa*

*Supplementary Table 6:* Summary statistics for *Y. aloifolia* and *Y. filamentosa* genes associated with CAM, N-metabolism, photorespiration, and citrate metabolism.

## Acknowledgements

The authors would like to thank Jen Liddle and Jeremy Balsbaugh for consultation on proteomic analysis, Kerrie Berry for project management at JGI, and Amir Ahkami and Sarah Leichty for project management at EMSL. We would also like to thank the Department of Energy Joint Genome Institute and collaborators for prepublication use of the genome and annotations for *Y. aloifolia* and *Y. filamentosa* for transcriptome analysis.

## Author Contributions

R.F., J.L-M, and K.H designed the research; R.F., K.W., R.C., J.T., N.T., N.M.M., H.H., E.S., and A.L. contributed to data collection, including physiology, transcriptomic, metabolomic, and proteomic; D.W. led analyses with assistance from R.F. and K.H.; D.W. wrote the paper with assistance from K.H.

## Conflict of interest

The authors declare no conflicts of interest.

## Funding

A portion of this research was performed under the Facilities Integrating Collaborations for User Science (FICUS) program (proposal:10.46936/fics.proj.2020.51552/60000215) and used resources at the DOE Joint Genome Institute (https://ror.org/04xm1d337) and the Environmental Molecular Sciences Laboratory (https://ror.org/04rc0xn13), which are DOE Office of Science User Facilities operated under Contract Nos. DE-AC02-05CH11231 (JGI) and DE-AC05-76RL01830 (EMSL).

## Data Availability

RNA seq data for our gene expression profiling analyses can be found in NCBI’s Sequencing Read Archive; SRA BioProject numbers: ’PRJNA1162856’, ’PRJNA1162850’, ’PRJNA1162828’, ’PRJNA1162846’, ’PRJNA1162829’, ’PRJNA1162847’, ’PRJNA1162862’, ’PRJNA1162861’, ’PRJNA1162852’, ’PRJNA1162845’, ’PRJNA1162853’, ’PRJNA1162857’. All other data, including physiological measurements, metabolite abundance, protein abundance, and normalized expression data are available on Dryad; DOI: 10.5061/dryad.5mkkwh7ft.

## Notes

https://github.com/dawickell/Yucca-omics

### Competing Interest Statement

The authors have declared no competing interest.

### Summary of Updates

Methods updated, version numbers added; Results pertaining to transcriptomic data reduced and simplified; citations and additional figure references added throughout; minor revisions to Figures 1,3,4,5; Sections re-ordered

