## Supplementary figures and images for "Integrative analysis of CAM photosynthesis reveals its impact on primary metabolism in *Yucca*"

### Supplementary Figures

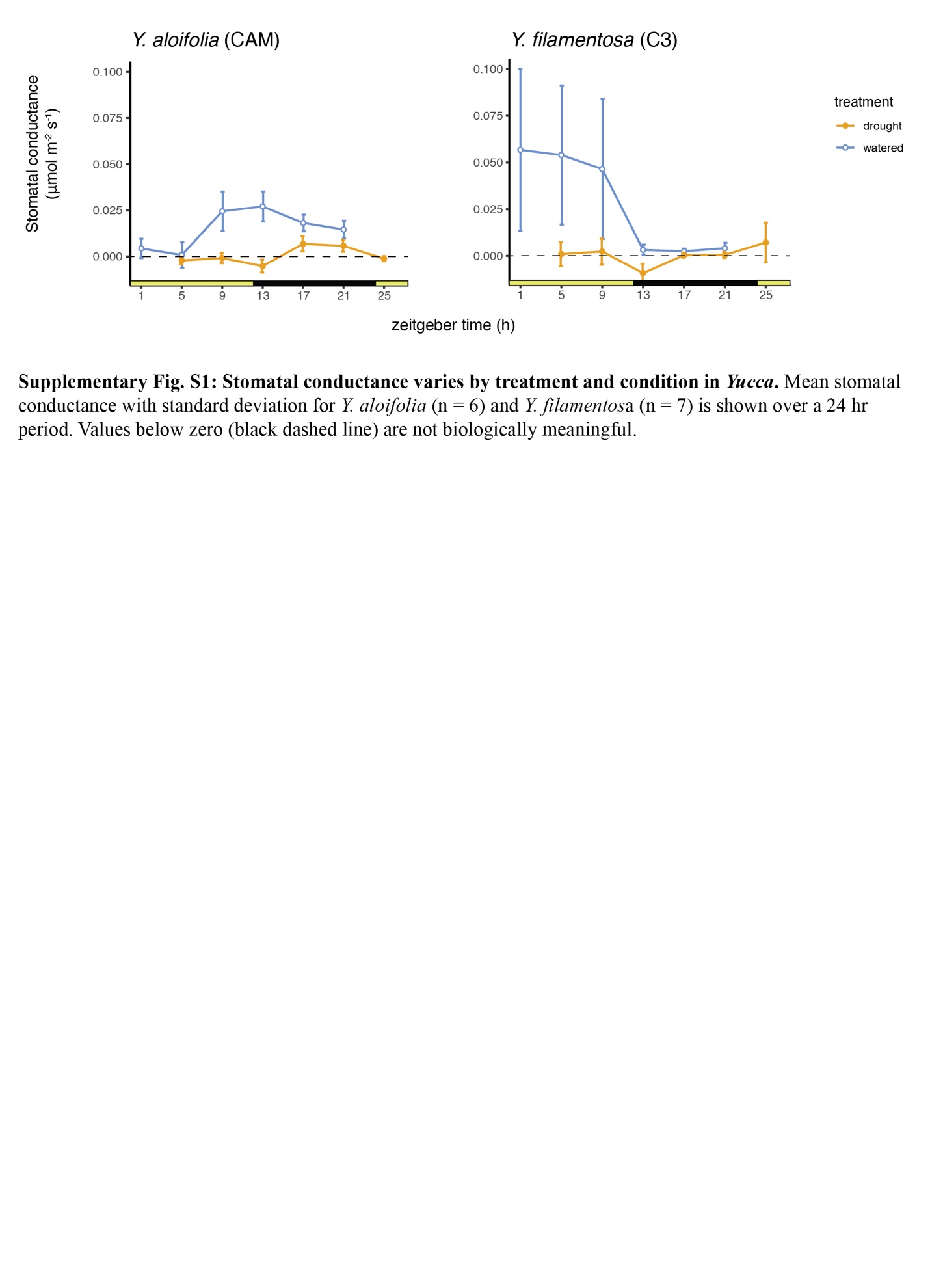

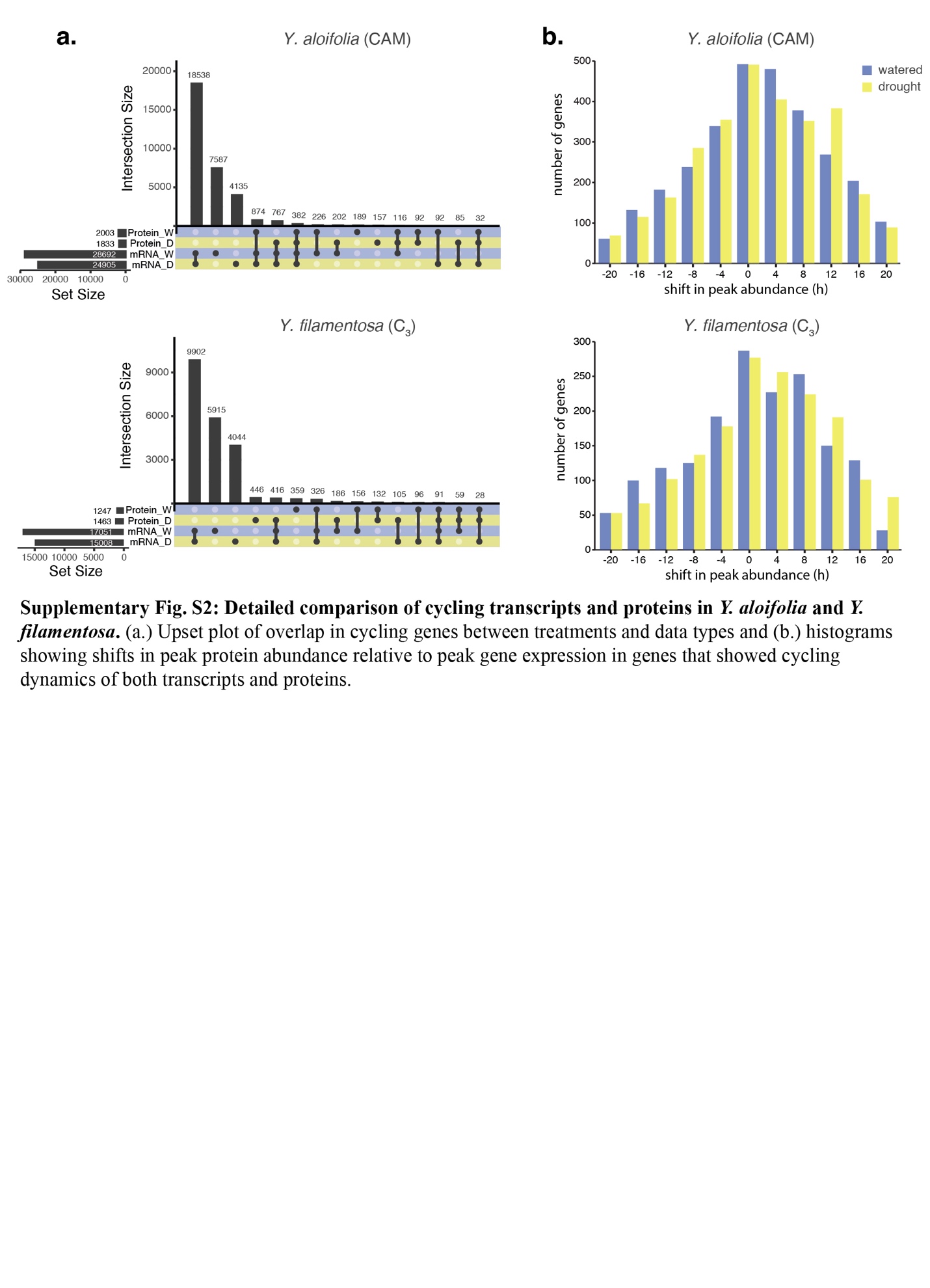

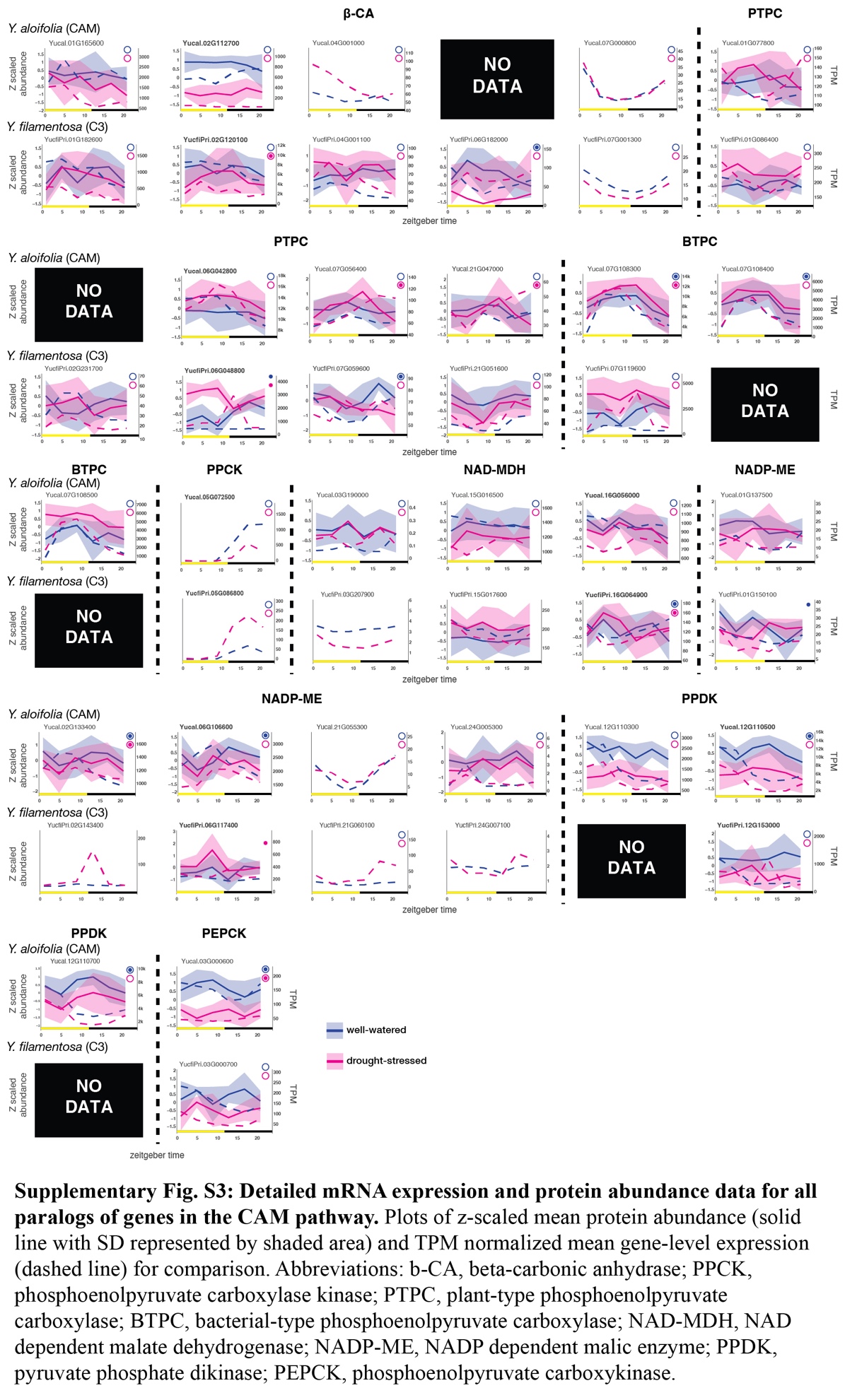

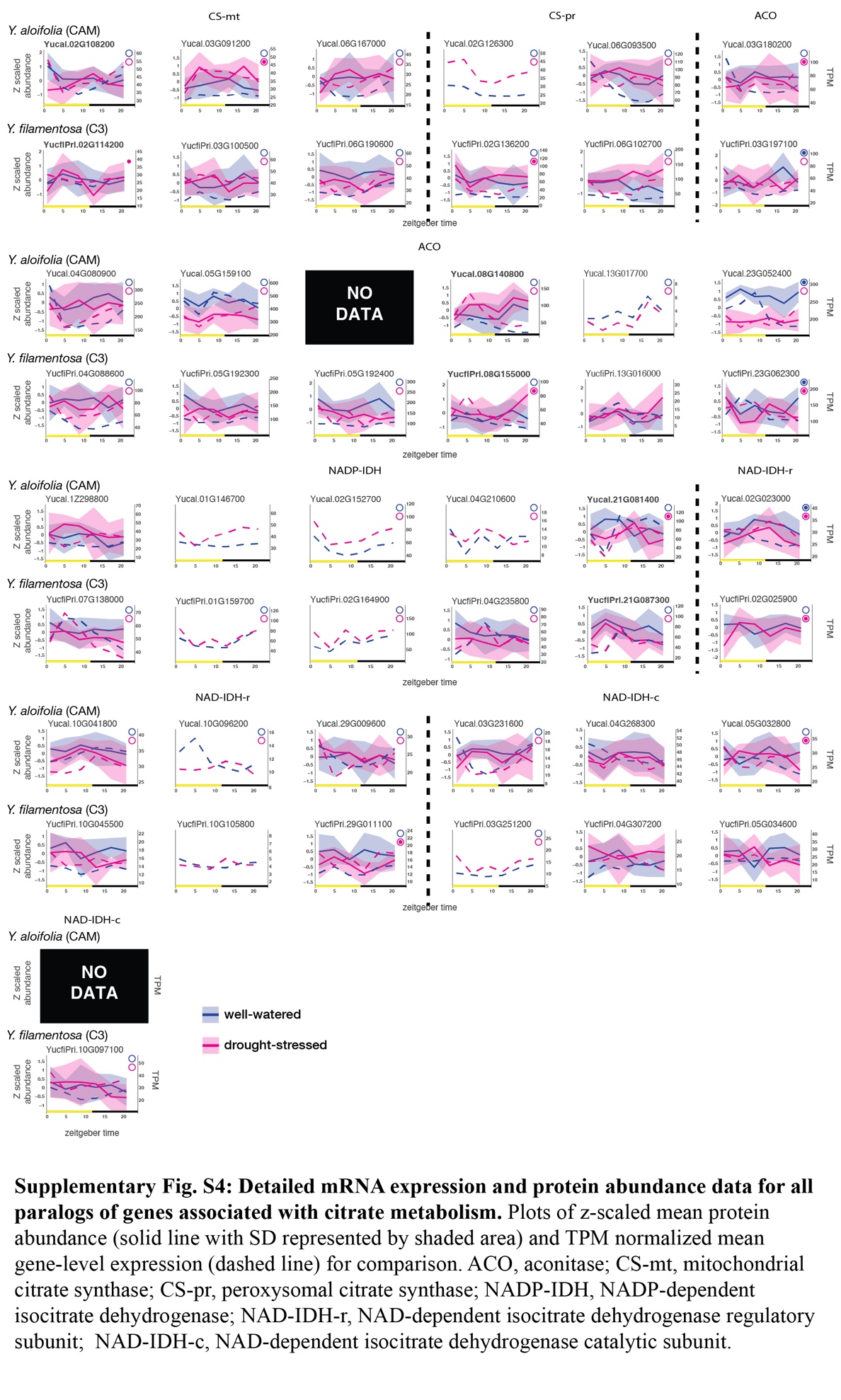

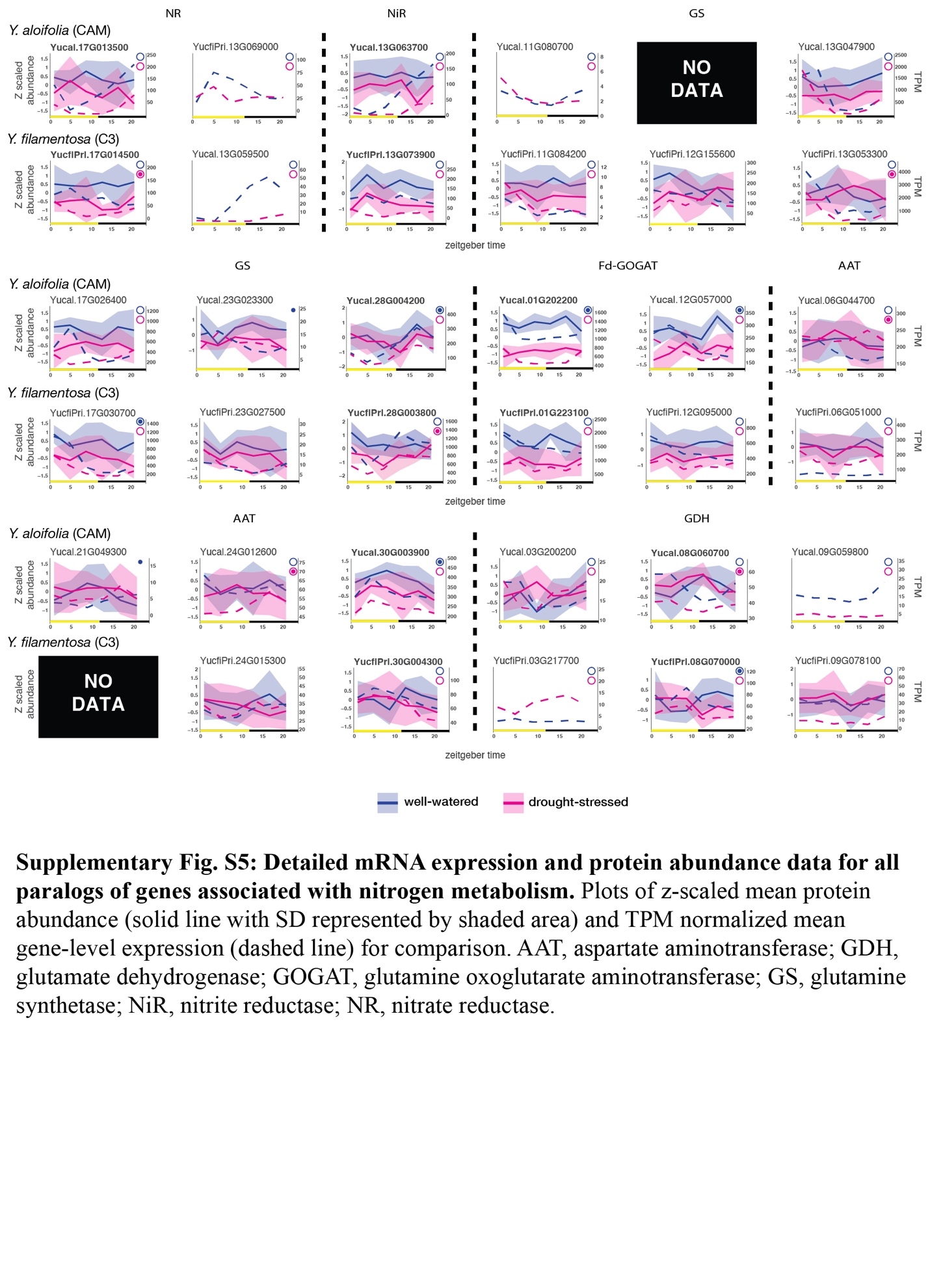

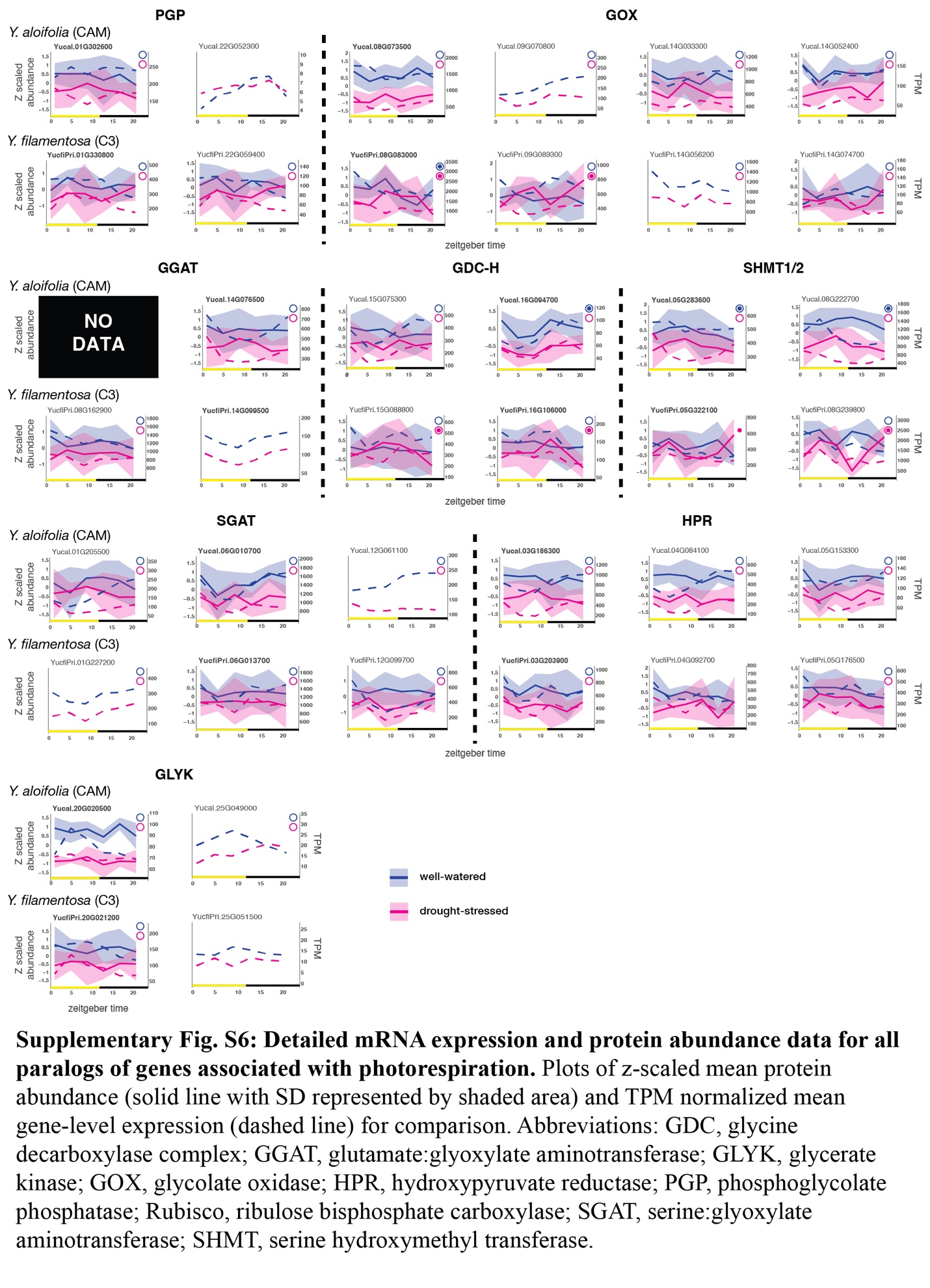
